# Combining segmental bulk- and single-cell RNA-sequencing to define the chondrocyte gene expression signature in the murine knee joint

**DOI:** 10.1101/2020.06.13.148056

**Authors:** Vikram Sunkara, Gitta A. Heinz, Frederik F. Heinrich, Pawel Durek, Ali Mobasheri, Mir-Farzin Mashreghi, Annemarie Lang

## Abstract

**Objective:** Due to the small size of the murine knee joint, extracting the chondrocyte transcriptome from articular cartilage (AC) is a major technical challenge. In this study, we demonstrate a new and pragmatic approach of combining bulk RNA-sequencing (RNA-seq) and single cell (sc)RNA-seq to address this problem.

**Design:** We propose a new cutting strategy of the murine femur which produces three segments with a predictable mixed cell populations, where one segment contains AC and growth plate (GP) chondrocytes, another contains GP chondrocytes, and the last segment contains only bone and bone marrow. We analysed the bulk RNA-seq of the different segments to find common and distinct genes between the segments. Then, the segment containing AC chondrocytes was digested and analysed via scRNA-seq.

**Results:** Differential expression analysis using bulk RNA-seq identified 350 candidate chondrocyte gene in the AC segment. Gene set enrichment analysis of these genes revealed biological processes related- and non-related to chondrocytes, including, cartilage development (adj. p-value: 3.45E-17) and endochondral bone growth (adj. p-value 1.22E-4), respectively. ScRNA-seq of the AC segment found a cluster of 131 cells containing mainly chondrocytes. This cluster had 759 differentially expressed genes which enriched for extracellular matrix organisation (adj. p-value 7.76E-40) and other joint development processes. The intersection of the gene sets of bulk- and scRNA-seq contained 75 genes, where all but ten genes were previously implicated in cartilage homeostasis or osteoarthritis (OA) progression.

**Conclusions:** Our approach has the potential to detect the scarce disease phenotypes of chondrocytes in murine OA models.

## Introduction

Knee osteoarthritis (KOA) is one of the top five disabling conditions that affects more than one-third of people aged over 65 years and over 100 million individuals globally [1]. KOA is characterized by an erosion of the articular cartilage, which leads to inflammation and pain accompanied by joint stiffness and immobility. Despite its prevalence, the molecular mechanisms triggering KOA are poorly characterized and current therapies cannot halt or revert KOA. While animal models of KOA exist, the causative mechanisms leading to cartilage erosion are poorly understood. It is therefore of utmost importance to identify the key molecular factors and the associated signalling pathways responsible for changes in chondrocyte function and the pathological transformation of healthy cartilage during the establishment of KOA [2–4]. Such molecular factors can serve as predictive biomarkers, and also represent targets for the therapy of KOA.

Preclinical models of KOA in mice are widely used to study the underlying molecular mechanisms of the disease [5, 6]. However, the dissection of articular cartilage (AC) from mice knee joints is challenging due to the small size, low thickness, and the huge potential for contamination with e.g. subchondral bone or surrounding connective tissue [7]. These technical challenges create obstacles for studying the gene expression signatures of AC chondrocytes. In addition, chondrocytes are isolated within a voluminous extracellular matrix (ECM) that is neither vascularised nor innervated and constitute a very small fraction of the whole knee joint [8]. Thus, attempts to identify the chondrocyte specific gene expression signatures hidden within the mixed transcriptome of various cell types using RNA-sequencing on population level (bulk RNA-seq) is almost impossible. A solution for this problem would be single cell (sc) RNA-seq of AC chondrocytes. However, the method requires the preparation of a single cell suspension from the tissue of interest, during which the potential influence of enzymatic, mechanical or chemical pre-treatment’s effects on the chondrocytes gene expression signature *in vivo* cannot be fully excluded [9, 10]. As chondrocytes are embedded in an ECM, an enzymatic digestion step is required. Hence, scRNA-seq analysis of musculoskeletal tissue such as bone and cartilage is a major challenge that has not been sufficiently resolved.

Here we present a new and pragmatic approach for extracting the gene expression signatures of chondrocytes from the articular cartilage (AC) of the murine knee joint using a combination of bulk RNA- and scRNA-seq. First, we propose a new cutting strategy for the joint which produces several control segments to help discriminate the chondrocyte-gene expression signature from the remaining joint cells’ transcriptome using bulk RNA-seq. Second, we describe for the first time a fast protocol for the digestion of AC and the isolation of chondrocytes followed by scRNA-seq. This protocol allows for an unaltered gene expression signature of individual chondrocytes *ex vivo* by using the transcriptional inhibitor actinomycin-D (Act-D) during digestion of the ECM. The combination of scRNA-seq and segmental bulk RNAseq enables a complete determination of the gene expression signature of murine AC chondrocytes, which will be key to understand underlying mechanisms leading to the establishment of KOA.

## Methods

Some sections of the methods, including histology, bulk RNA library preparation and sequencing, bulk RNA-seq alignment and normalization, scRNA-seq alignment, normalization, clustering and differential expression analysis as well as gene set enrichment analysis are further described in the Supplementary methods.

### Animals and tissue collection

Male C57BL/6J mice aged 10 weeks (n = 6) were purchased from Charles River Laboratories and were euthanized at an age of 12 weeks by cervical dislocation. The knee joints were collected, and the femur was carefully dissected. The femur was cut in 3 defined segments – segment A included the articular surface, subchondral bone and a small growth plate area of the femur (lateral and medial condyles). Segment B contained growth plate, bone marrow, non-loaded articular cartilage (*Facies patellaris femoris*) and bone while segment comprised bone marrow and bone. The cutting procedure was verified via histology and weight analysis.

### Bulk RNA isolation

For bulk RNA isolation, segments were transferred to RNAlater (Qiagen) for 1 h at 4°C before removing the RNAlater and freezing samples at −80°C. Segment samples were cryo-pulverized (59012N, Biospec) and gently resuspended in TriFast™ (VWR), mixed with 1-bromo-3-chloropropane (Sigma Aldrich) and incubated for 10 min. Centrifugation was performed for 10 min at 10.000 *x g* and the top aqueous phase was further collected for RNA isolation using the RNeasy Mini Kit (Qiagen) according to the manufactures’ instructions. Purity of the RNA was analyzed via Nanodrop; RNA integrity and quality were verified via Agilent 2100 Bioanalyzer or Fragment Analyzer (RNA-sequencing).

### Tissue digestion and cell preparation for single-cell RNA-sequencing

Femur preparation was done on ice with PBS supplemented with 2 μg/mL Actinomycin D (Sigma Aldrich). Segments were digested in 10 mg/ml Collagenase II (Nordmark) supplemented with 2 μg/mL Actinomycin D for 4 h at 37°C and constant rotation. The cell suspension was transferred to a cell strainer and washed with PBS/BSA. FACS analysis (MacsQuant, Miltenyi Biotec) was performed to determine cell concentration and vitality (DAPI staining; > 80 % cell vitality as cut off).

### Single cell RNA-sequencing

For single cell library preparation, single cells were applied to the 10X Genomics platform using the Chromium Single Cell 5’ Library & Gel Bead Kit (10× Genomics) and following the manufacturer’s instructions for capturing ~3,000 cells. The amplified cDNA was used for 5’ gene expression (GEX) library preparation by fragmentation, adapter ligation and index PCR. The quality of final single cell 5’ GEX libraries was assessed by Qubit quantification and fragment analysis (DNF-474 High Sensitivity NGS Fragment Analyzer Kit, Agilent). The sequencing was performed on a NextSeq500 device (Illumina) using High Output v2 Kits (150 cycles) with the recommended sequencing conditions for 5’ GEX libraries (read1: 26nt, read2: 98nt, index1: 8nt, index2: n.a.).

## Results

### Cutting of the murine femur provides sufficient RNA per segment and predictable mixed cell populations in each segment

Chondrocytes can be found in the articular cartilage (AC) and the growth plate (GP) of the knee joint (Fig. 1a). We isolated the femur from mice and dissected it into three segments, labelled A, B and C. Segment A included the medial and lateral femoral condyle up to the GP; while segment B comprised the GP and metaphyseal area; and lastly, segment C contained half of the diaphysis (Fig. 1b). Given the small size of the murine joint, it was not possible to cut the femur condyles without including small parts of the GP. For this reason, the diaphyseal segment C served as a control for the signal interference from the GP in segment A and B. The cutting protocol was verified using H&E and Safranin-O staining, which clearly outlined the AC and GP in segment A, and the GP in B. There was no cartilage present in segment C (Fig. 1c). The cutting protocol also produced comparable samples for each segment, giving an average signal to noise ratio in their wet weight of approximately 4.6 (N=6), meaning there was no more than 21% variation in the cutting within each segment (Fig. 1d). Upon isolation of the RNA from each segment, we found that all samples yielded more than 1,000 ng of total RNA (Fig. 1e), with segment A yielding the lowest amount of RNA among the segments in the range of 2,772 ng to 6,135 ng. The RNA integrity number (RIN) values of the segment samples were found to be of sufficient quality for sequencing, with all segments showing a RIN value above 8 (Fig. 1f). In summary, each mouse joint was cut into three consistent segments with a predictable mixed cell population, and each segment contained adequate RNA quantity and quality for RNA-seq analysis.

**Figure 1:**
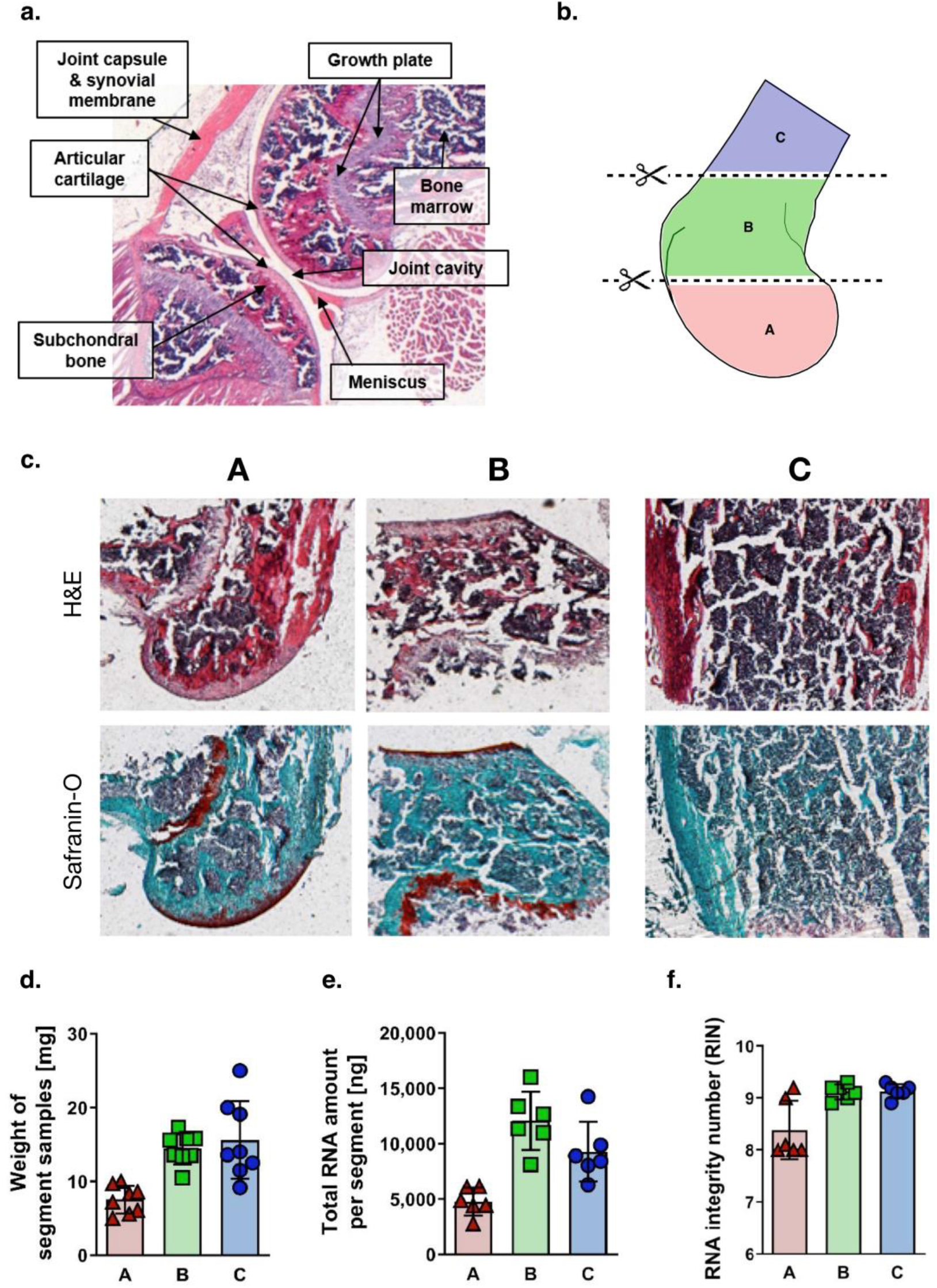
The proposed segments of the mouse joint contain sufficient RNA per segment and predictable mixed cell populations. **(a)** H&E staining of sagittal cross-section of the murine knee joint. **(b)** Schematic indicating the approximate cutting region of the three segments of the femur. **(c)** Staining of sagittal cross-sections of representative segments: A, B, and C. H&E staining (top), Safranin-O staining (bottom): cartilage is red coloured. Scale bars indicate 200 μm. **(d)** Weight of samples of the segments represented as a bar graph (mean ± STD) and individual data points (n= 8). **(e)** Total RNA amount per segment represented as a bar graph (mean ± STD) and individual data points (n= 6). **(f)** RNA integrity number per segment represented as a bar graph (mean ± STD) and individual data points (n= 6).

### Bulk RNA-seq analysis of the segments reveal AC chondrocyte candidate genes but cannot guarantee their chondrocyte specificity

Since we aimed at delineating the transcriptome of AC chondrocytes and to reduce the signal interference from GP chondrocytes, we performed differential gene expression analysis between segment A and C, segment B and C, and segment A and B (Fig. 2a–c). To identify the differentially expressed genes, a Mann-Whitney U-test with a 3-fold change cut-off and a p-value of 0.05 was used. We found 350 genes which were upregulated in segment A with respect to segment C. Gene set enrichment analysis (GSEA) found these genes enriched in biological processes involved in cartilage development (GO:0051216, adj. p-value: 3.54E-17, genes: 28) and chondrocyte development (GO:0002063, p-value: 1.56E-4, genes: 8) (Fig. 2e). A comparable amount—274 genes—was upregulated in segment B when compared to segment C. These genes are involved in similar biological processes as those found in the comparison between segment A and C, such as cartilage development. However, there were also genes enriched in growth plate cartilage development (GO:0003417,adj. p-value: 8.81E-4, genes: 6) and endochondral bone growth (GO:0003416, adj. p-value:1.22E-4, genes: 7) (Fig. 2f). As a next step, we studied the genes differentially expressed between segment A (containing AC and GP chondrocytes) and segment B (containing only GP chondrocytes) and found 71 genes to be upregulated in segment A. These we assumed to be potentially AC chondrocyte-specific. However, GSEA of these genes did not show enrichment in any biological process (Fig. 2d).

**Figure 2:**
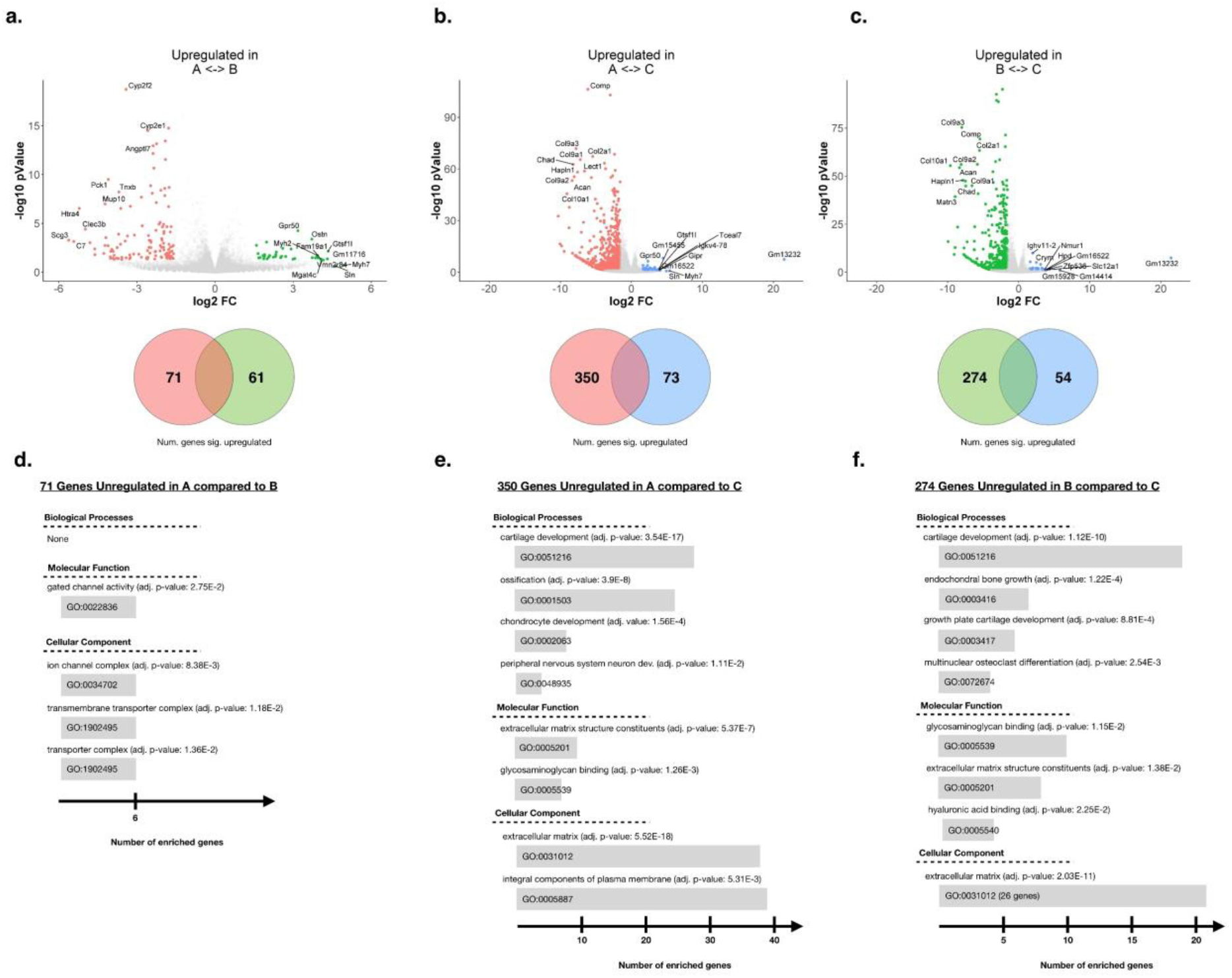
Bulk RNA-seq analysis of the segments reveal potential AC-specific genes. **(a–c)** Volcano plots (top) highlighting the top ten genes differentially expressed between the two segments of interest. Venn diagram (bottom) stating the total number of genes which showed a larger than 3-fold change difference in expression between the two segments of interest. (n= 3) **(a)** Comparison between segment A and segment B. **(b)** Comparison between segment A and segment C. **(c)** Comparison between segment B and segment C. **(d–e)** Biological processes, molecular functions, and cellular components found in the gene set enrichment analysis (GSEA) of the gene set of interest. The associated GO terms are given in a bar graph with the length of the bar indicating the number of genes found in that term. **(d)** GSEA of genes more than 3-fold upregulated in segment A compared to B. **(e)** GSEA of genes more than 3-fold upregulated in segment A compared to C. **(f)** GSEA of genes more than 3-fold upregulated in segment B compared to C.

Other than chondrocyte-related processes, we also found that genes upregulated in segment A with respect to segment C were involved in the biological processes of peripheral nervous system neuron development (GO:0048935, adj. p-value: 1.11E-2, genes: 4) and ossification (GO:0001503, adj. p-value: 3.91E-8, genes: 25) (Fig. 2e). Similarly, among genes upregulated in segment B compared to segment C, we found genes enriched in growth plate cartilage development (GO:0003417, adj. p-value: 8.81E-4, genes: 6) and multinuclear osteoclast differentiation (GO:0072674, adj. p-value: 2.54E-3, genes: 4). This suggests that the segments contain genes from cells other than chondrocytes, for example, pre-osteoblasts, nerve cells and osteoclasts.

In summary, by juxtaposing the transcriptomes of the segments A, B, (containing chondrocytes) and C (containing no chondrocytes), we found a high variety of genes that may be AC chondrocyte-specific. However, we were not able to discern the transcriptomes of the different cell populations contained in the different segments.

### scRNA-seq analysis of the condylar segment A reveals that chondrocyte-like cells are scarce but cluster as a distinct population

Since we assumed both chondrocyte phenotypes–from AC and GP–to be present in segment A, three samples were separately digested and analysed via scRNA-seq. Because we could not see any sequencing batch effects (Fig. 3a), the data from the three different experiments were combined for analysis. The aggregation of the three batches led to a total of 7,133 cells to be analysed. The Seurat-3 analysis pipeline deduced six clusters of different cell subpopulations (Fig. 3b). Using one versus all differential gene expression analysis with a cut-off of 3-fold change and GSEA, we discerned the cell subpopulations per cluster (see Supplementary Results A). Cluster 1 to 5 were enriched for hematopoietic stem cells. Therefore, cluster 1, 3, 4, and 5 comprised myeloid cells while cluster 2 included the lymphoid cells (Fig. 3). In more detail, cluster 1 was enriched for granulocytes (N=2,073) while cluster 4 comprised mainly neutrophils (N=789) which was also supported by the fact that both clusters showed overlapping gene expression (Supplementary Results A). Cluster 5 contained monocytes and macrophages (N=317) and showed expression of genes unique for antigen presenting cells which was comparable with cluster 2 (B-cells and precursors; N=1,336) (Fig. 3c; Supplementary Results A). Cells from the erythroid lineage and precursors were found in cluster 3 (N=1,055) which was supported by the identification of the corresponding biological processes (Fig. 3c). In cluster 6 mainly chondrocytes were found, even though the presence of osteoblast-specific genes indicated the presence of other musculoskeletal cells and GP chondrocytes (N=131).

**Figure 3:**
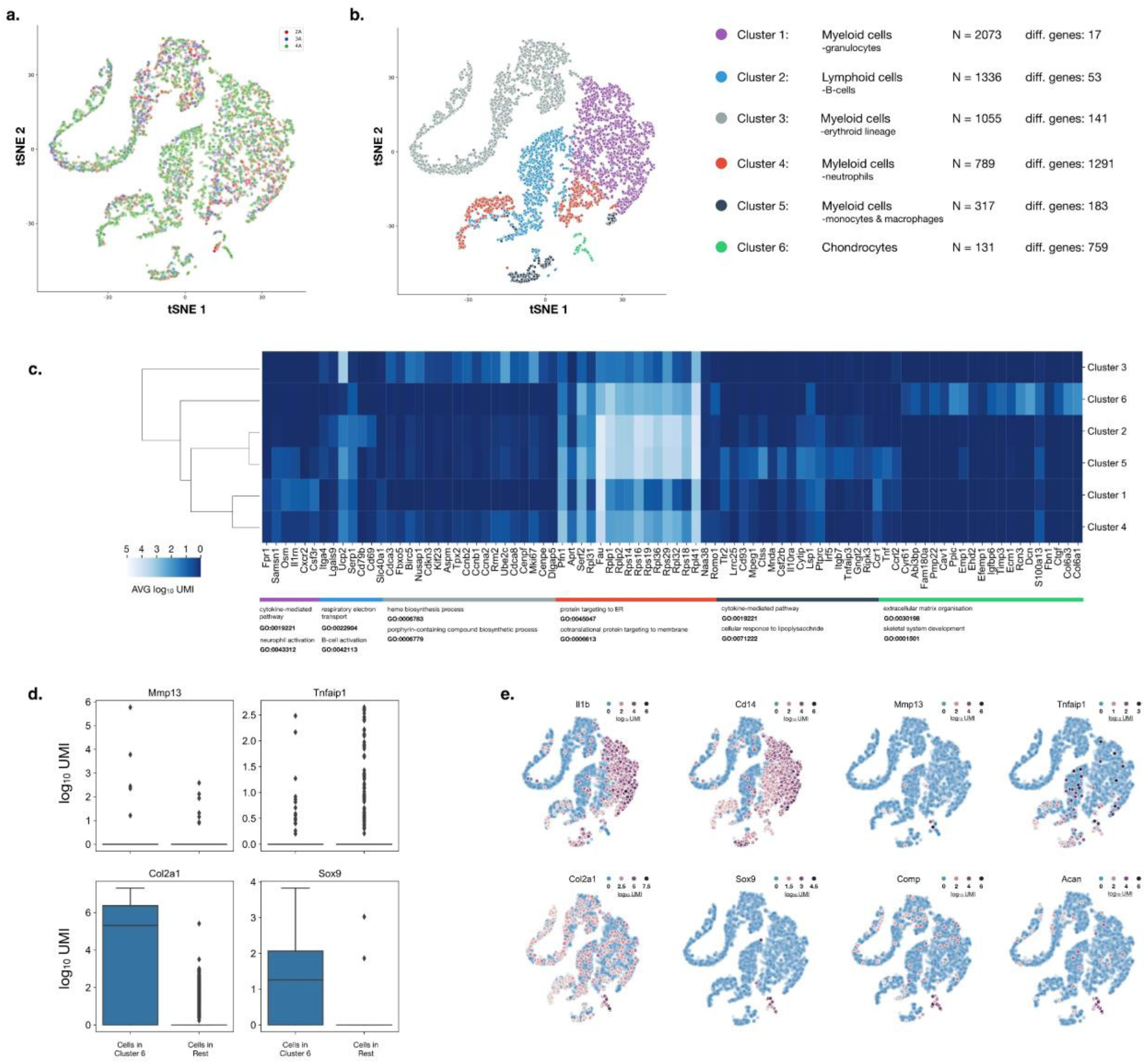
scRNA-seq analysis of the condylar segment A show chondrocyte-like cells cluster. **(a)** tSNE plot of the scRNA-seq analysis from segment A, with the cell colour highlighting the batch to which the cell belongs. **(b)** tSNE plot highlighting the cluster to which cells have been allocated and a list of the potential cell phenotypes in the clusters, the number of cells belonging to each cluster, and the number of differentially expressing genes (3-fold change cut-off) in comparison to all other clusters. **(c)** Columns of the heatmap contain selected differentially expressed genes in each of the clusters, rows contain their expression of in each cluster. Dendrogram showing the hierarchy of the clusters over the selected genes. **(d)** Box plot showing the expression of *Mmp13*, *Tnfaip1, Col2a1,* and *Sox9* of cells within and outside cluster 6. **(e)** tSNE plots highlighting the magnitude of expression of the genes of interest in the cells from the scRNA-seq assay of segment A.

In order to identify the chondrocytes in cluster 6, we searched for cells expressing cartilage-specific genes such as *Sox9* (89 cells; 68%), *Col2a1* (94 cells; 72%), and both genes together (78 cells; 60%) (Fig 3d, 3e). We concluded that the 78 cells expressing both *Sox9* and *Col2a1* were chondrocytes from AC or GP. Next, we asked the question whether the transcriptome of these cells could be affected by the digestion protocol in order to exclude false positive results. Since the digestion bias has not yet been addressed in detail in the sequencing field, we specifically looked for well-known effects of digestion. As a results, we did not observe an upregulation of *Mmp1*, *Mmp13*, *Tnfaip1*, *ColX*, and a downregulation of *Col2A1*, *Col1A2*, *Acan* (see Supplementary Results B). Hence, the addition of ActD during tissue digestion seemed to preserve the phenotypic identity of the chondrocytes and appeared to inhibit potential digestion-induced alterations in the transcriptome.

GSEA of the 759 genes differentially expressed in cluster 6 (containing 60% chondrocytes) found some of these genes to be enriched in processes involving cartilage development (GO:0051216, adj. p-value: 1.65E-27, genes: 57), bone development (GO:0060348, adj. p-value: 7.3E-17, genes: 42), nervous system development (GEO:0007399, adj, p-value: 3.21E-10, genes: 143), and angiogenesis (GEO:0001525, adj. p-value:6.83E-12, genes: 57). Thus, we could deduce that the cells in cluster 6 are involved in production and upkeep of the ECM and bone in the joint (Fig. 4), and that they are a mixed population of chondrocytes, osteoblasts, and other cells.

**Figure 4:**
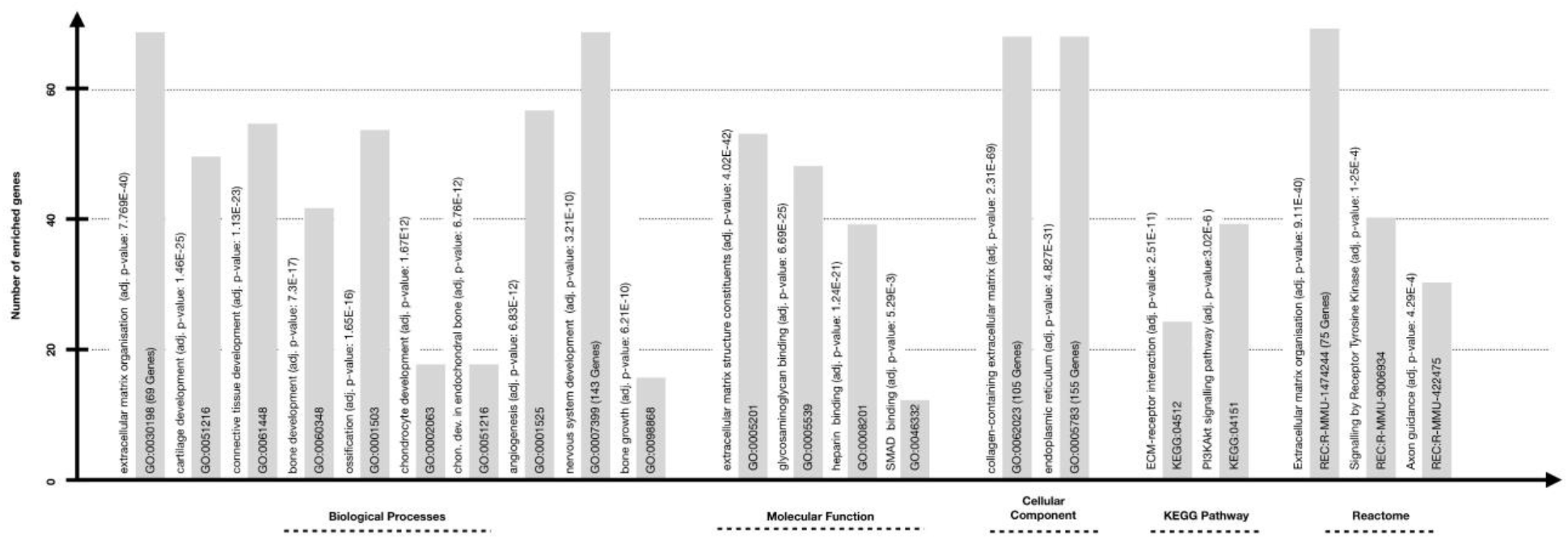
Gene set enrichment analysis of the 788 genes found to be differentially expressed in cluster 6 compared to all other clusters. Results originate from enrichment searches in the KEGG pathway, Reactome pathway, and Gene Ontology (Biological Processes, Molecular Function, and Cellular component) databases. The associated terms are given in a bar graph with the length of the bar indicating the number of genes found associated with that term. The name of the term and the adjusted p-value are stated adjacent to the bar on the left side.

Using scRNA-seq, we were able to extract a large set of genes belonging to the chondrocyte transcriptome. However, due to the small population size of the AC chondrocytes, they could not be discriminated from bone and other synovium-associated cells.

### Both approaches yield complementary data sets

To date, there is no complete transcriptomic atlas of the cells in the murine joint. The well accepted assumption is that the transcriptome of the AC chondrocytes and GP chondrocytes have a larger overlap. If we use binary classification nomenclature, and say the genes resulting from the analysis of the respective methods found AC chondrocyte “positive” genes, and the remaining genes to be AC chondrocyte “negative” genes, then we can see that both approaches identify different sets of genes to be AC chondrocyte positive (see Fig. 5a for illustration). From our GSEA, we saw that the AC chondrocyte positive gene-set from bulk RNA-seq’s also contained genes well-known to be expressed in nerve cells. These genes, in binary classification nomenclature, are “false positive” AC chondrocyte genes. In contrast, in scRNA-seq, we observed the “false positive” genes to be genes associated with GP chondrocytes. Interestingly, the two approaches identified different, mutually exclusive, “false positive” genes. Thus, we hypothesised that using the intersection of the genes detected by both approaches would eliminate the “false positive” genes and enable us to delineate the AC chondrocyte transcriptome.

**Figure 5:**
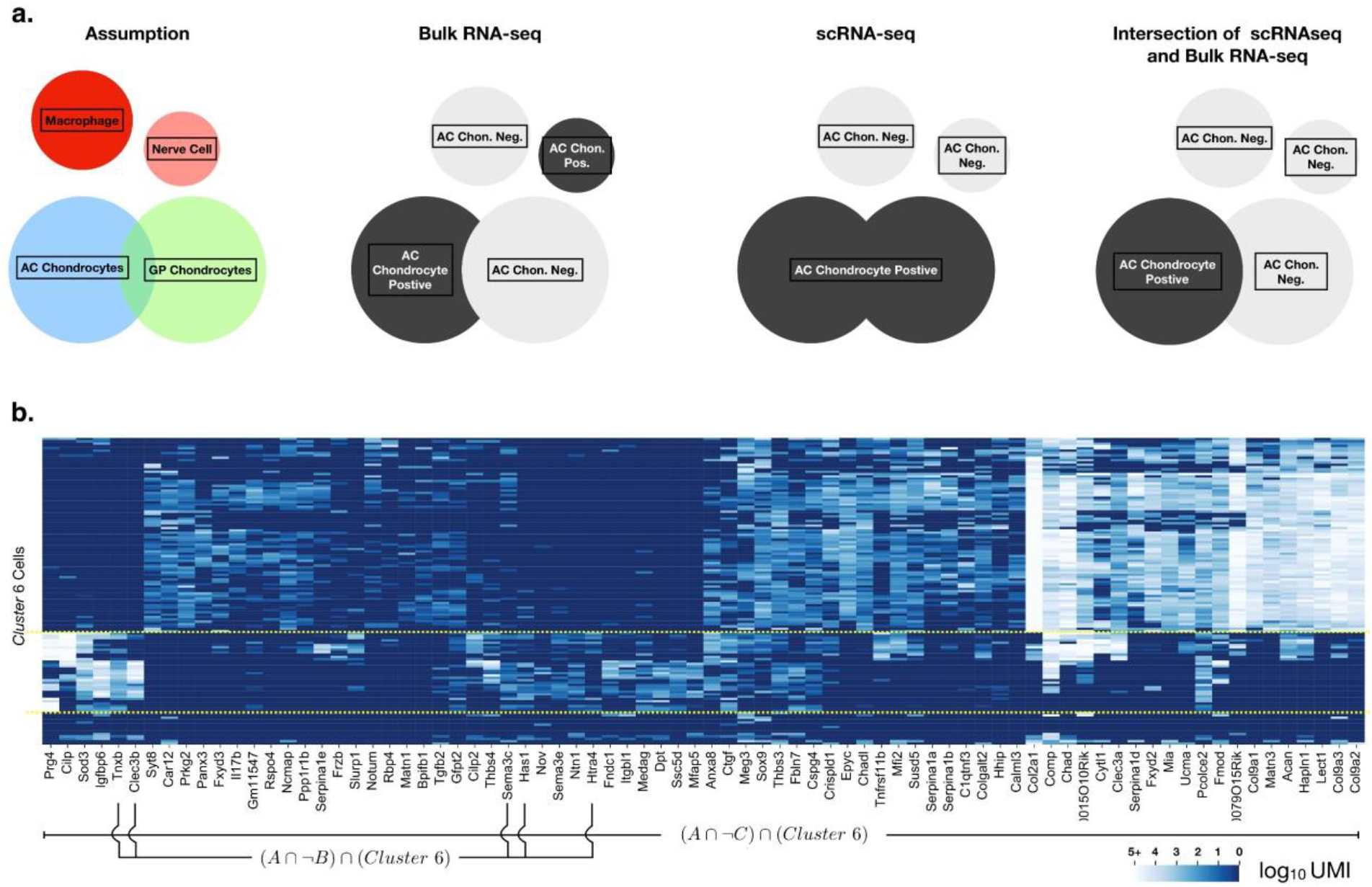
RNA-seq and scRNA-seq identify different, false positive genes. **(a)** Schematic of Venn diagrams. Each circle represents the unique gene set of the cell type given in the label. Classification of the cell types in the left figure as chondrocyte positive or negative by technique stated in the caption. From left to right, bulk RNA-seq misclassifies genes belonging to nerve cells as belonging to chondrocytes, and likewise, scRNA-seq misclassifies genes belonging to bone related cells as belonging to chondrocytes. The intersection of both methods results in only the true chondrocyte positive gene set remaining. **(b)** Columns show the genes contained in the intersection of differentially expressing genes of cluster 6 (scRNA-seq) and the differentially expressing genes of segment A compared to segment C and segment A compared to segment B. The rows contain the cells allocated to cluster 6 (scRNA-seq). Each square of the heat map represents the expression of the gene (column) in the cell (row).

### Using the intersection of bulk RNA-seq and scRNA-seq yields the AC chondrocyte-specific transcriptome

To build a robust gene set of AC chondrocytes from the femoral cartilage, we combined all genes upregulated in segment A with respect to segment C and the genes upregulated in segment A with respect to segment B from the bulk RNA-seq. This gene set presumably contained chondrocyte-related genes. Then, we further reduced this gene set to chondrocyte-specific genes by taking the intersection of the genes in this set with genes upregulated in cluster 6 from the scRNA-seq data of segment A. This intersection gene set contained 75 genes (Fig. 5b, see Supplementary Results C: Table 1). In depth literature search revealed that a considerable number of these genes were AC cartilage-specific and had already been studied in the context of cartilage homeostasis or OA pathogenesis. The majority of genes encode for matrix proteins such as *Matn1* and *3*, *Lect1*, *Chadl*, *Epyc*, *Chad*, *Col9*, *Col2a1*, *Acan*, *Comp*, *Fmod*, *Ucma*, *Cilp2*, *Hapln*, *Clec3a* and *3b*, *Tnbs3* and *4*, *Fndc1*, *Prg4*, *Dpt*, *Cspg4* or factors and enzymes contributing to matrix formation or degradation: *Serpina 1a*, *1b*, *1d* and *1e*, *Prkg2*, *Car12*, *Tnxb*, *Htra4*, *Pcolce2*, *Gfpt2*, *Colgalt2*, *Has1* and *Ppp1r1b*. Out of the 75 genes, there were only 10 genes which could not be related to cartilage or a specific function, these were *Slurp1, Calml2, Ssc5d, Crispld1, Bpifb1, Mfap5, Medag, Gm11547, 1500015O10Rik* and *3110079O15Rik*. Additionally, genes that are related to cellular functions were detected. For example, *Sod3* is abundant in the cartilage ECM maintaining the homeostasis by controlling reactive oxygen species (ROS). *Nov/Ccn3* encodes a matricellular protein with relevance in chondrocyte metabolism that represses endochondral ossification. *Mfi2*, *Syt8*, *Panx3*, *Anxa8*, *Fxyd 2* and *3*, *Itgbl1* and *Ntn1* are related to the active transport, cell membrane fusion, integrin interaction, or encode for transmembrane proteins. Some of these genes have been described to act anabolic (*Mfi2*, *Syt8*, *Fxyd2* and *3*, *Itgbl1*) while the rest is pro-catabolically supporting the progression of OA (*Panx3*, *Anxa8*, *Ntn1*). Moreover, inflammation-associated genes such as *Il17b, Cytl1, C1qtnf3, Rbp4* seemed to be expressed even under homeostatic conditions maintaining the balance between anabolic and catabolic factors. Interestingly, several genes were identified that are related to specific signalling pathways, for example, Wnt signalling (*Notum, Tnfrsf11b, Frzb, Rspo4*), indian hedgehog signalling (*Hhip*), Sox9 signalling (*Mia*), Smad signalling (*Meg3*), support cartilage differentiation, and maturation as growth factors (inhibitors) such as, *Sox9, Ctgf*, *Tgfb2*, and *Igfbp6*. *Sema 3c* and *3e* and *Ncmap* were ranked as genes with association to the nervous system relevant for axonal guidance and myelin formation, respectively.

**Table 1:**
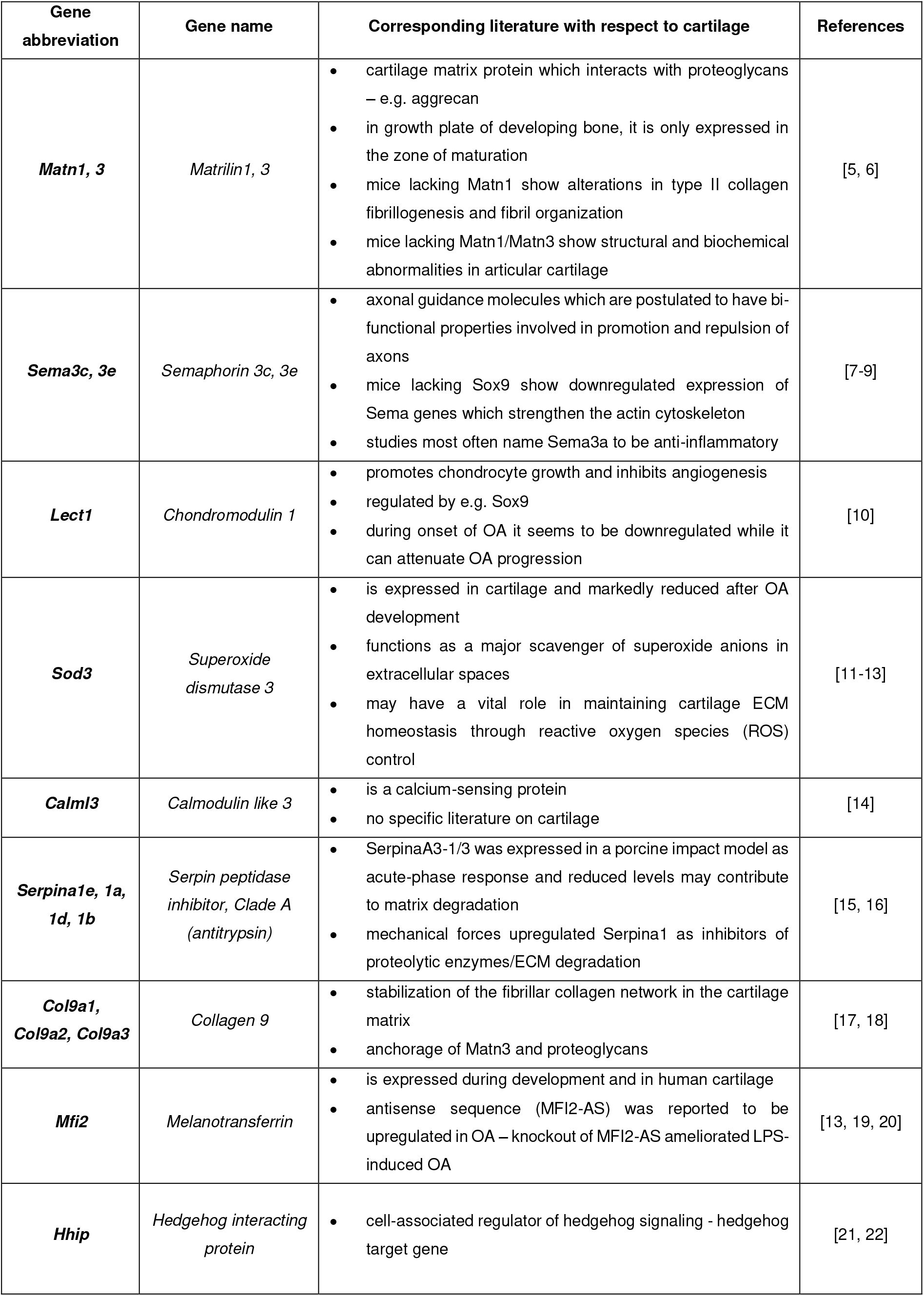
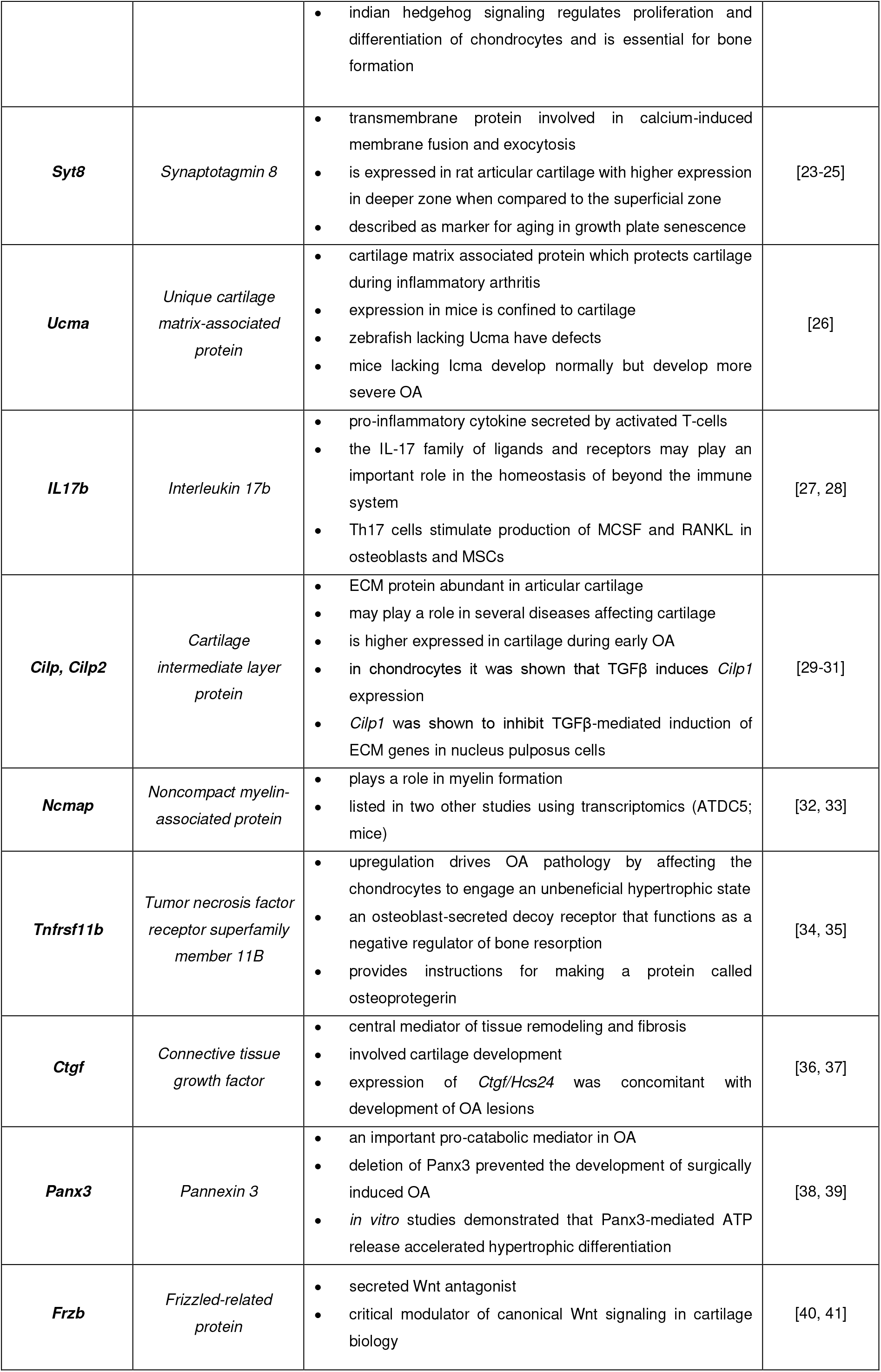
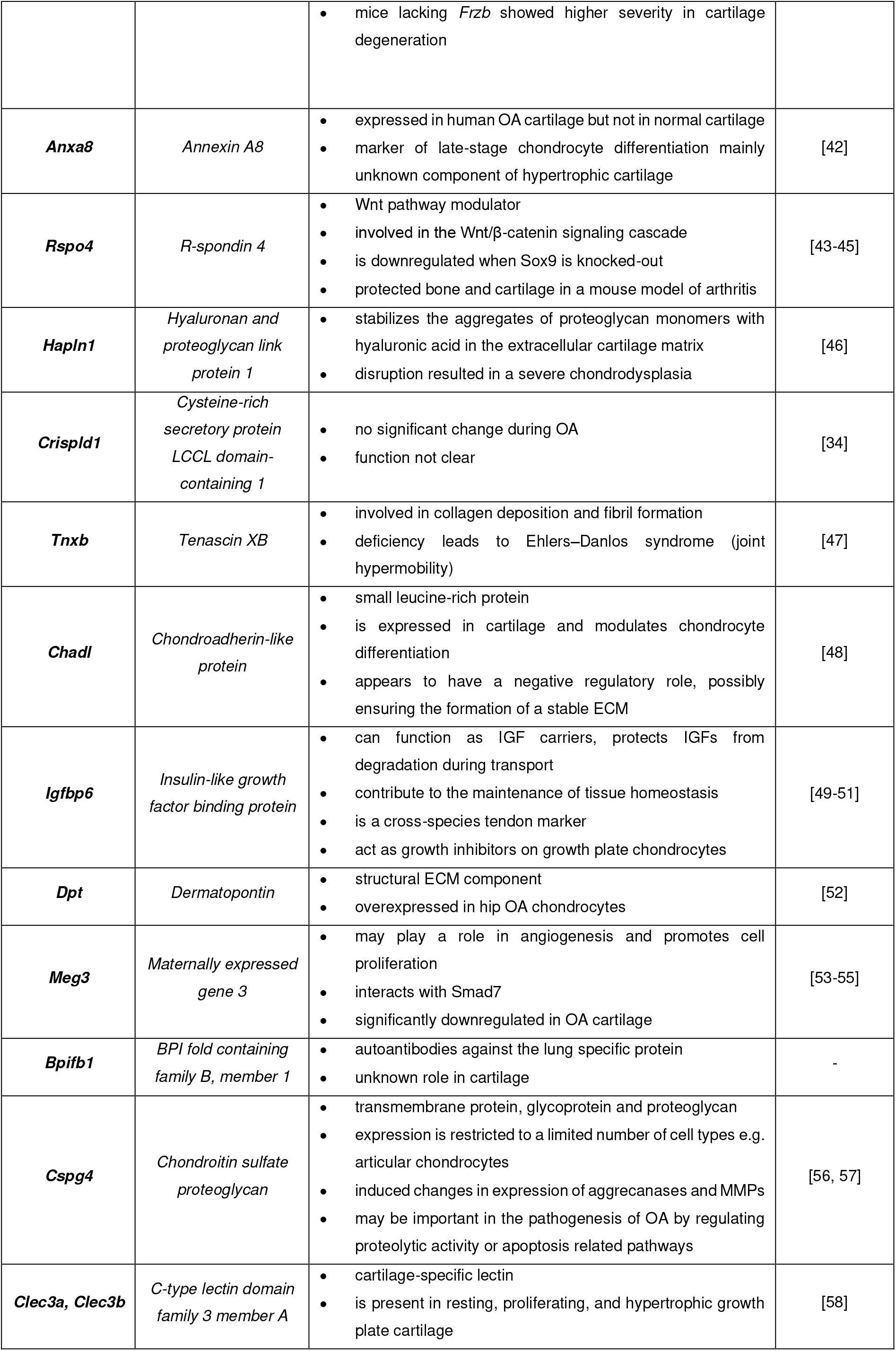
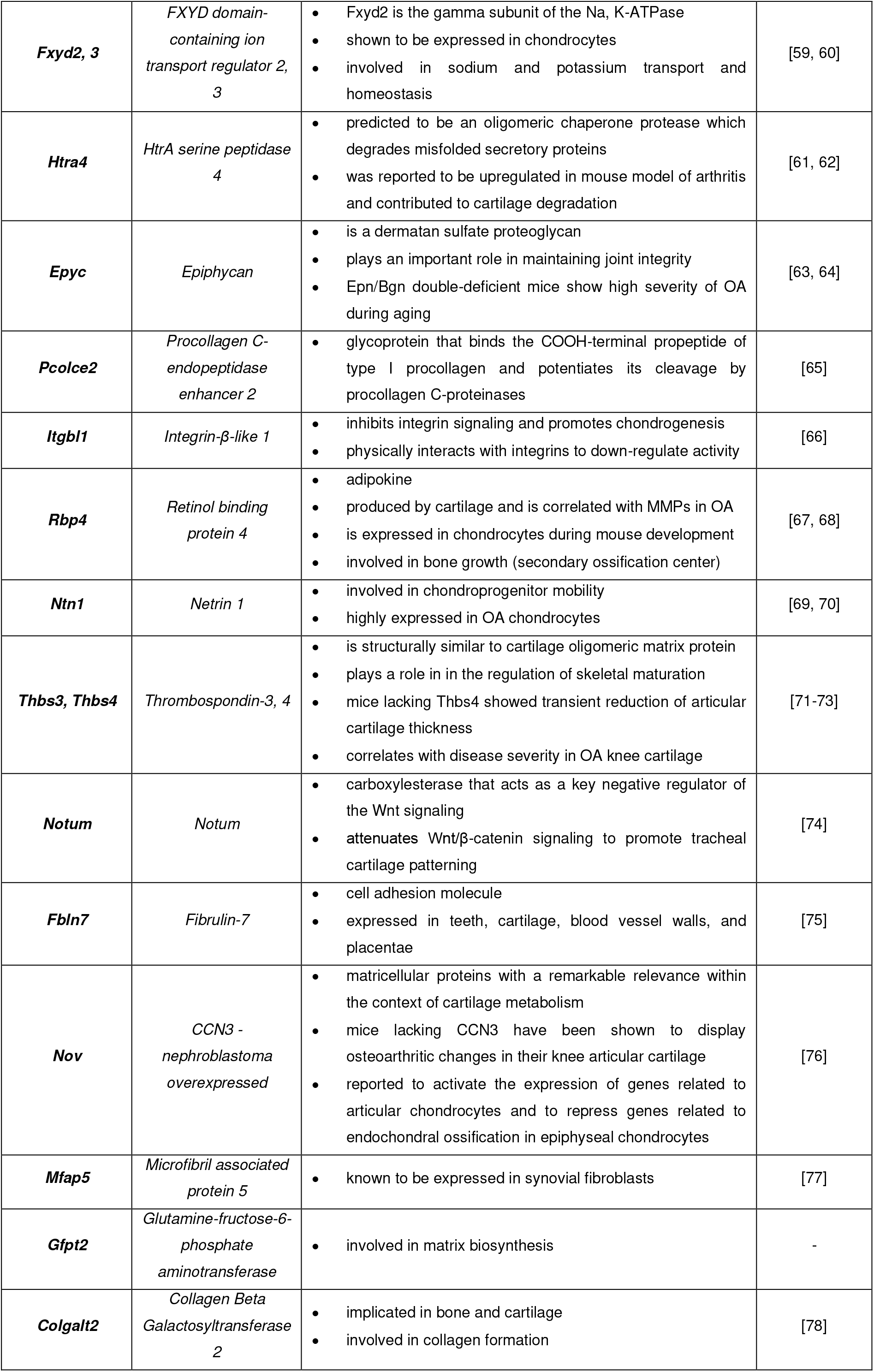
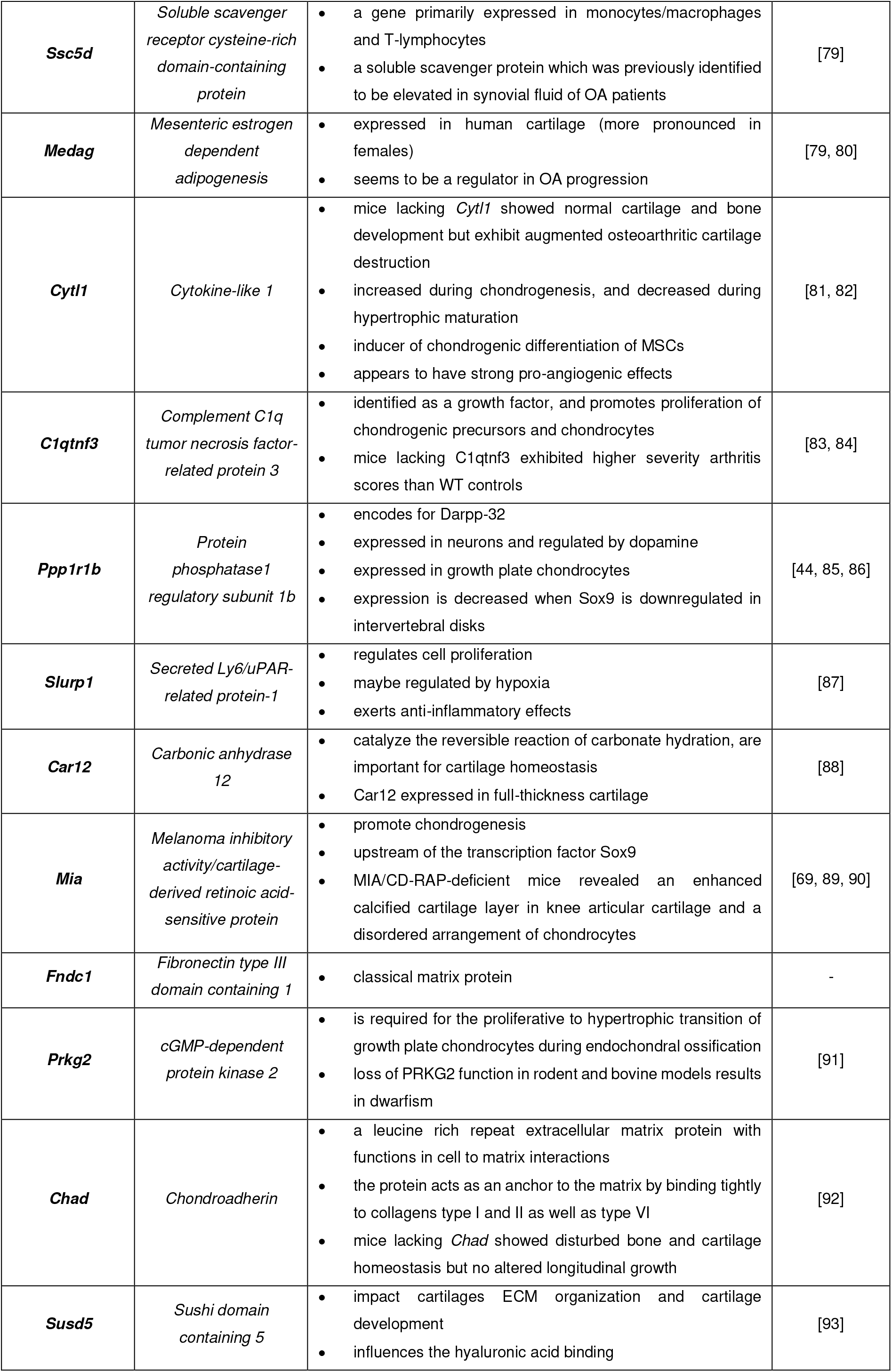
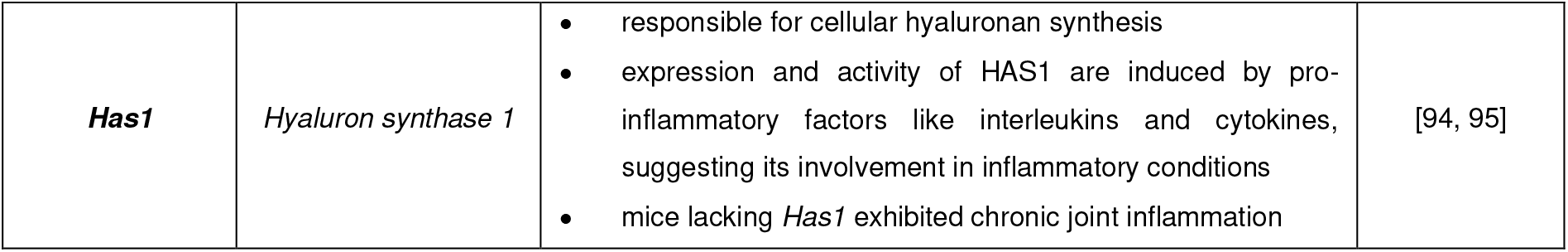
Selected differentially expressed genes which are AC chondrocyte-specific. -well-known genes such as *Col2a1, Acan, Sox9, Tgfb2, Comp, Fmod, Prg4* are not additionally listed

## Discussion

The only cell type that can be found in fully developed mature articular cartilage is the chondrocyte, with a low species-specific volume density of 2 % (human) and 15–40 % (mouse) [11, 12]. The overall cartilage thickness on the femoral ends in the knee joint is reported to be approximately 2.35 mm in humans and 0.1 mm in mice [11, 13, 14]. It was found by Gardiner et al., that cartilage only contributes less than 0.2 % of RNA to the total RNA in the murine knee joint [7]. Due to this low resolution, they proposed to compare pathways rather than genes when performing meta-analysis of RNA studies on joint tissues. Though this is a step in the right direction, the approach still cannot exclude that many genes and pathways that have been described so far are attributed to potential other cell and tissue contamination within the samples. To overcome these unresolved technical issues, we proposed the novel and pragmatic approach of using multiple tissue segments–each segment with at least one mutually exclusive phenotype in its population–to be used as controls to delineate phenotypes of mixed cells in the joint. In particular, to delineate the AC chondrocyte transcriptome in the knee joint, we cut three segments, one segment containing AC and GP chondrocytes, one segment with only GP chondrocytes, and the last segment containing no chondrocytes. Using these three segments, we were able to delineate genes upregulated in AC chondrocytes and GP chondrocytes. The cutting strategy gave a robust gene-set list for chondrocytes; however, the approach could only capture the upregulated genes and not the whole chondrocyte transcriptome.

To date, insights into the single cell transcriptome of murine chondrocytes can only be gathered from limb developmental studies, where different stages in chondrocyte phenotype progression/development are described, for example, the formation of the growth plate [15, 16] or different bone marrow niches [17]. Hence, the literature on the chondrocyte single cell transcriptome in the murine joint is vacuous. In our study, scRNA-seq analysis of the condylar segment A revealed that chondrocyte-like cell cluster distinctly in the population but suffer from scarce cell presence. Nevertheless, we found that these chondrocyte-like cells had hundreds of genes which are significantly upregulated with respect to other cells in the joint, hence, giving them a distinct phenotypic signature. Due to lack of surface markers for cell sorting, we could only extract approx. 78 chondrocytes (1.1 % of all cells analysed) out of three femoral condyles. Even though the proportion of chondrocytes extracted seems low, it is in fact consistent with the proportion of chondrocytes which are found in the joint [7]. Taken together, the chondrocyte transcriptome is distinct enough to be detected in single cell assays, however, due to the small sampling frequency, heterogeneity in chondrocytes could not be detected with confidence.

Due to the nature of the problem of isolating chondrocytes in murine joints, we found that both bulk RNA-seq and scRNA-seq do suffer from poor specificity. Hence, we suggest performing both methods and using the intersecting genes as the true chondrocyte candidate genes. In this work, the intersection of bulk RNA-seq and scRNA-seq yielded 75 genes which were AC chondrocyte-specific. Interestingly, beside the classical chondrocyte markers which are mainly related to ECM formation, degradation and integrity, we found new genes which were previously not associated with chondrocytes. For example, the gene *Sema3*, which has been described in context of axon promotion and repulsion [18] and was also found to be transactivated by *Sox9* [19] Similarly, *Ncmap* plays a role in myelin formation and has also been listed in other transcriptome studies [7, 20]. Presumably, these genes are mainly expressed in deep layer chondrocytes of the osteochondral unit and mediate peripheral nerve growth. However, their implication on cartilage homeostasis, OA progression or even pain sensitization is unknown. In addition, *Clec3a* and *b* encode a cartilage-specific lectin which has been shown to be upregulated in GP chondrocytes but has not been described in detail in articular cartilage [21]. Vice versa, *Htra4*, a oligomeric chaperone protease which degrades misfolded secretory proteins has only been described in the context of arthritis and its involvement in cartilage degradation [22, 23]. Further genes such as *Slurp1, Calml3, Ssc5d, Crispld1, Bpifb1, Mfap5* and *Medag* were not related to cartilage or surrounding tissue at all, providing directions for further research studies. For examples, *Slurp1* regulates cell proliferation and exerts anti-inflammatory effects [24], while *Ssc5d* has been reported to be expressed in monocytes/macrophages and T-lymphocytes [25] and *Mfap5* can be found in synovial fibroblasts [26].

To isolate chondrocytes from cartilage, the treatment with collagenases is indispensable. Dissociation-induced alterations of the transcriptome are widely discussed in the community, although not very often studied or included in publications [10]. Notably, there is a fine line between the concentration of the collagenase, the digestion time, and the cell output, which need to be adjusted precisely to keep negative influences on the chondrocyte transcriptome as low as possible [9, 27, 28]. It was suggested that an additional treatment with transcription inhibitors might resolve this problem [29, 30]. Hence, in our study, we supplemented Actinomycin D (ActD), a chemotherapeutic medication and transcription inhibitor which has been shown to freeze the transcriptome in immune cells [31]. We did not find collagenase-induced upregulation of genes, suggesting that the ActD treatment paused the digestion-induced transcriptional changes. It must be noted that the chondrocytes in this experiment were healthy. In an OA (disease) setting, the chondrocytes are known to be under stress and undergoing differentiation or apoptosis. Future research is needed to investigate if ActD can preserve chondrocytes which are in a transient phase during disease progression.

Regarding the technical aspects, the natural next step is to achieve a higher yield of chondrocytes from the murine joint for scRNA-seq analysis. Due to the lack of surface markers, this has been proven to be a challenge. However, in our experimental workflow we were able to highlight the different cell populations appearing in scRNA-seq of the murine joint. Hence, we can use our results to tag all cells which are not chondrocytes, resulting in an up-sampling of chondrocyte and chondrocyte-like cells. For example, we can use CD14 to exclude monocytes/macrophages, Ter119 to exclude erythrocytes/erythroblasts, and CD34 to exclude bone marrow cells.

We believe that our approach of combining the two RNA-seq methods will be effective in studying the disease phenotypes of chondrocytes in murine OA model in the future. Thus, target genes could be identified supporting the development of chondrocyte phenotype-specific treatment strategies.

## Acknowledgement

We like to thank Prof. Frank Buttgereit, Dr. Timo Gaber and Dr. Max von Kleist for scientific discussions and support.

## Contributions

V.S., M.F.M. and A.L. designed the study. V.S., A.L. and G.A.H. collected and analyzed data. V.S., F.H. and P.D. performed analysis of bulk RNA-seq and scRNAseq data. V.S. and A.L. prepared the main manuscript text. M.F.M., G.A.H. and A.M. revised manuscript. All authors finally approved the manuscript.

## Supplementary Methods

### Histology

For histology femur segment samples were fixed with 4% paraformaldehyde at 4°C for 6 h and ran through a sucrose gradient (10 %, 20 %, 30 %) for 12-24h each. Cryo-embedding was performed using SCEM medium (Sectionlab) and cryo-slices of 7 μm were produced using cryo-films (Sectionlab) and stored at −80°C.

Hematoxylin and Eosin (H&E) staining: air drying, fixation with 4% PFA (10 min), washing with Aqua dest., first staining step in Harris’s haematoxylin solution (7 min) (Merck, Germany), washing twice with Aqua dest., differentiation step in 0.25% HCl-solution, washing twice with tap water, second staining step in 0.2% Eosin (2 min) (Chroma Waldeck, Germany), differentiation and washing in 96% ethanol and 100% ethanol, fixation with xylol, covering of stained slices with Vitro-Clud® (R. Langenbrinck GmbH).

Safranin-O & light green staining: air drying, fixation with 4% PFA (10 min), washing with Aqua dest., first staining step in 1% Safranin-O solution (8 min) (Sigma Aldrich), washing twice with Aqua dest., second staining step in 0.1 % Light green (5 min) (Sigma Aldrich, Germany), washing in 1 % acetic acid and 100% ethanol, fixation with xylol, covering of stained slices with Vitro-Clud® (R. Langenbrinck GmbH).

### Bulk RNA library preparation and sequencing

The mRNA-specific library preparation was performed using 10 ng of total RNA for each sample. For poly-A dependent cDNA synthesis and a first amplification step the Smart-Seq v4 mRNA Ultra Low Input RNA Kit (Clontech) was used according to the manufacturer’s instructions. After quality control (DNF-474 High Sensitivity NGS Fragment Analyzer Kit, Agilent) and concentration measurement (Qubit dsDNA HS Assay Kit, Invitrogen), 1 ng of the purified cDNA was used for tagmentation and library completion with the Nextera XT library preparation kit (Illumina). Paired-end sequencing (2×75 nucleotides) was performed on a Illumina NextSeq500/550 using a Mid Output v2 Kit (150 cycles, Illumina).

### Bulk RNA-Seq alignment and normalization

NextSeq500 (Illumina) raw sequence reads were mapped to the mouse GRCm38/mm10 genome with TopHat [1] in very-sensitive settings for Bowtie2 [2]. Gene expression was quantified by featureCounts [3].

### Bulk RNA-Seq differential expression analysis

For the analysis of the mRNA and differentially expressed genes, DESeq2 [4] within R 3.5.1 in default settings was used.

### Single cell RNA-seq alignment and normalization

Illumina output was demultiplexed and mapped to the mm10 reference genome by cellranger‐3.0.2 (10× Genomics Inc.) using refdata‐cellranger‐mm10‐1.2.0 in default parameter setting and 3000 expected cells.

### Single cell RNA-seq clustering and differential expression analysis

Raw UMI‐counts were further analyzed using R 3.6.1 with Seurat package [5], as proposed by Butler and colleagues, including log‐normalization of UMI‐ counts, detection of variable genes and scaling. T‐distributed Stochastic Neighbour Embedding and the underlying Principle Component Analysis was performed based on 30 components using variable genes and a perplexity of 30 as set by default.

### Gene Set Enrichment Analysis

Gene set enrichment analysis was performed using g:profiler (version e99_eg46_p14_f929183). The default g:GOSt method was used for calculating significance scores with correction for multiple tests [6]. We choose our adjusted p-value cut-off to be 0.05. The databases: GO molecular function, GO cellular components, GO biological process, KEGG, and Reactome were used for the enrichment.

## Supplementary Results

### Supplementary Results A: Classification of the subpopulations identified with scRNA-seq in segment A

Subpopulations were clustered based on the expression of specific genes and the enrichment analysis of certain biological processes. Cluster 1 enriched for granulocytes as supported by the expression of i) *CD33*, a myeloid-specific marker, ii) *Cxcr2*, which is known to be prominent on neutrophils, iii) *CD300lf*, an inhibitory receptor for myeloid cells and mast cells and iV) *Csf3r*, the granulocyte colony-stimulating factor (see supplementary Figure A). This is in line the found biological processes such as cytokine-mediated pathway (GO:0019221) and neutrophil activation (GO:0043312). In addition, gene expression overlaps were found with cluster 4 (neutrophils) for e.g. *CD177*, the neutrophil antigen B, *Camp*, found in polymophonuclear leukocytes and *Ly6g*, a prominent marker for neutrophils. The biological processes such as protein targeting to ER (GO:0045047) and cotranslational protein targeting to membrane (GO:0006613) are not exclusively know for a specific cell type.

Cluster 2 shows high gene expression of *CD79a* and *b* as well as *Ms4a1* (CD20) specific markers for B-cells and B-cell precursors supported by the expression of CD69 an activation or retention marker. Biological processes enriched for respiratory electron transport (GO:0022904) and B-cell activation (GO:0042113).

Cells of the erythroid lineage can be clearly found in cluster 3 indicating the pronounced expression of genes encoding for enzymes involved in heme synthesis e.g. *Alas2* or protein unique for e.g. erythrocytes or erythroblasts e.g. *Hba-a1, Abcg* and *Hemgn*. Moreover, biological processes such as heme biosynthesis (GO:006783) and porphyrin-containing compound biosynthetic process (GO:0006779) are well known from erythroblasts and erythrocyte recycling [1].

In addition, cluster 5 enriched for monocytes and macrophages showing classical marker such as *CD14, CD74* and *Ccl2*. Furthermore, *Tnf* was expressed which is known to be upregulated during monopoesis. With respect to biological processes cytokine-mediated pathways (GO:0019221) and cellular response to lipopolysaccharide (GO:0071222) were identified.

Finally, cluster 6 enriches for mainly chondrocytes (*Prg4, Chad*) which is stringently supported by the identified biological processes: extracellular matrix organization (GO:0030198) and skeletal system development (GO:0001501). However, the presence of osteoblast-specific markers (*Col1, Sparc, Sp7*) indicated the presence of GP chondrocytes and other musculoskeletal cells.

**Supplementary Figure A:**
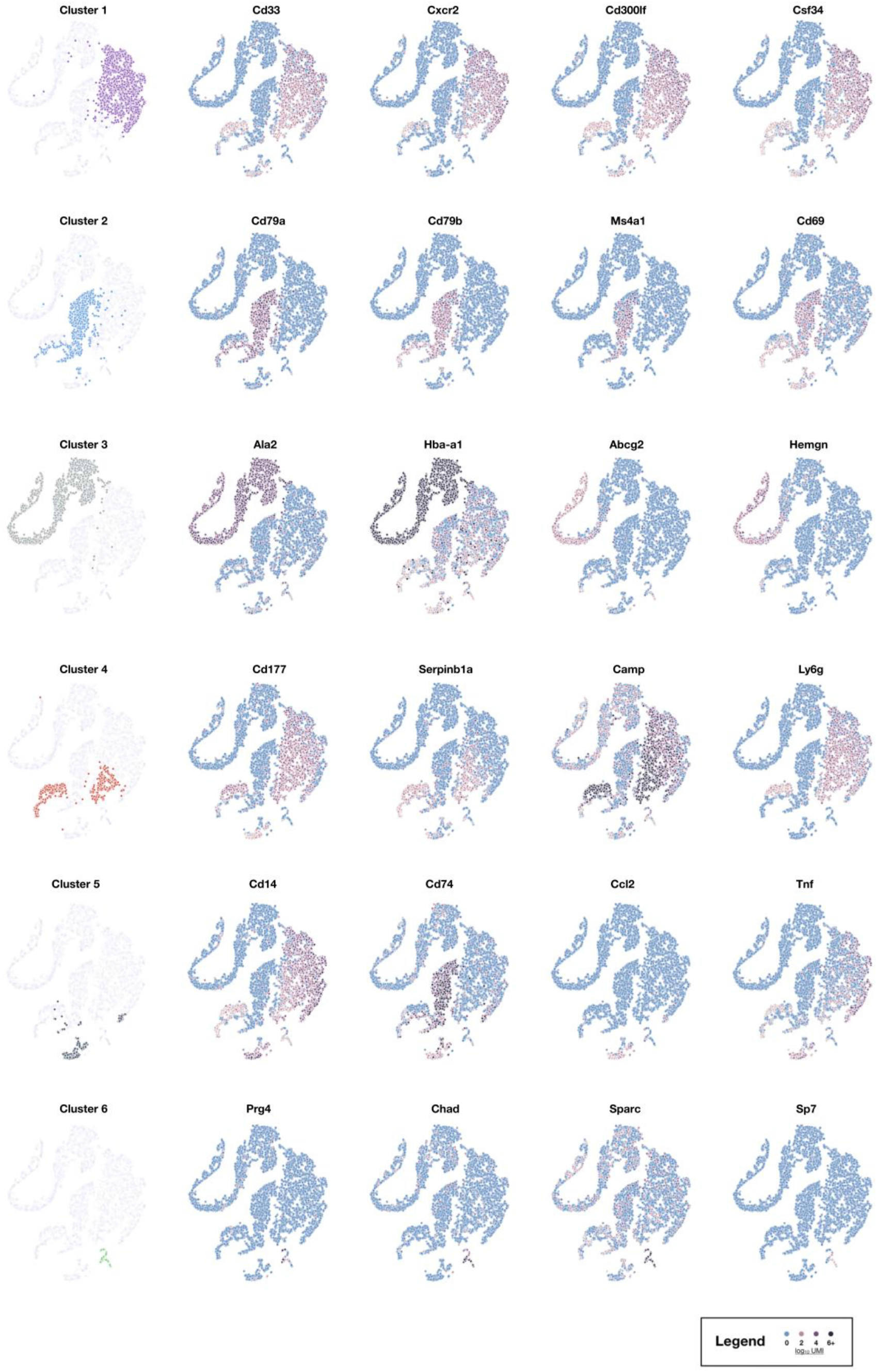
Classification of subpopulations. tSNE plots highlighting the cluster to which cells have been allocated and selected differentially expressing genes which support the allocation of distinct cell populations to the clusters.

### Supplementary Results B: Effects of Actinomycin D on the single cell transcriptome

We know from literature that *Sox9* is not modulated during the ECM digestion [2]. Hence, we isolated the transcriptome of the cells in cluster 6 which had a non-zero *Sox9* expression. Then, we normalized the transcriptome of the cells by the Sox9 expression and did the same with the transcriptome of the segments in bulk RNA-seq. Comparing the *Sox9* normalized expression levels of the cells in cluster 9 and the segments, we did not see an upregulation of genes *Mmp13, Tnfaip1, ColX* in the scRNA-seq compared to bulk (supp. Fig. B a–c). Similarly, we did not see the down regulation of *Col2a*, *Acan*, and *Comp* in the scRNA-seq compared to bulk (supp. Fig. B d–f). This suggests that the Act-D was successful in inhibiting at-least the previously establish digestion effects on chondrocytes.

Furthermore, we did find the upregulation of FAM46C encoding an active non-canonical poly(A) polymerase which has been described to support mRNA stability and gene expression [3]. Hu et al. reported the upregulation of FAM46C expression after ActD treatment [4]. The upregulation of this genes in our single cell samples indicates the effectiveness and functionality of the ActD treatment [3].

**Supplementary Figure B:**
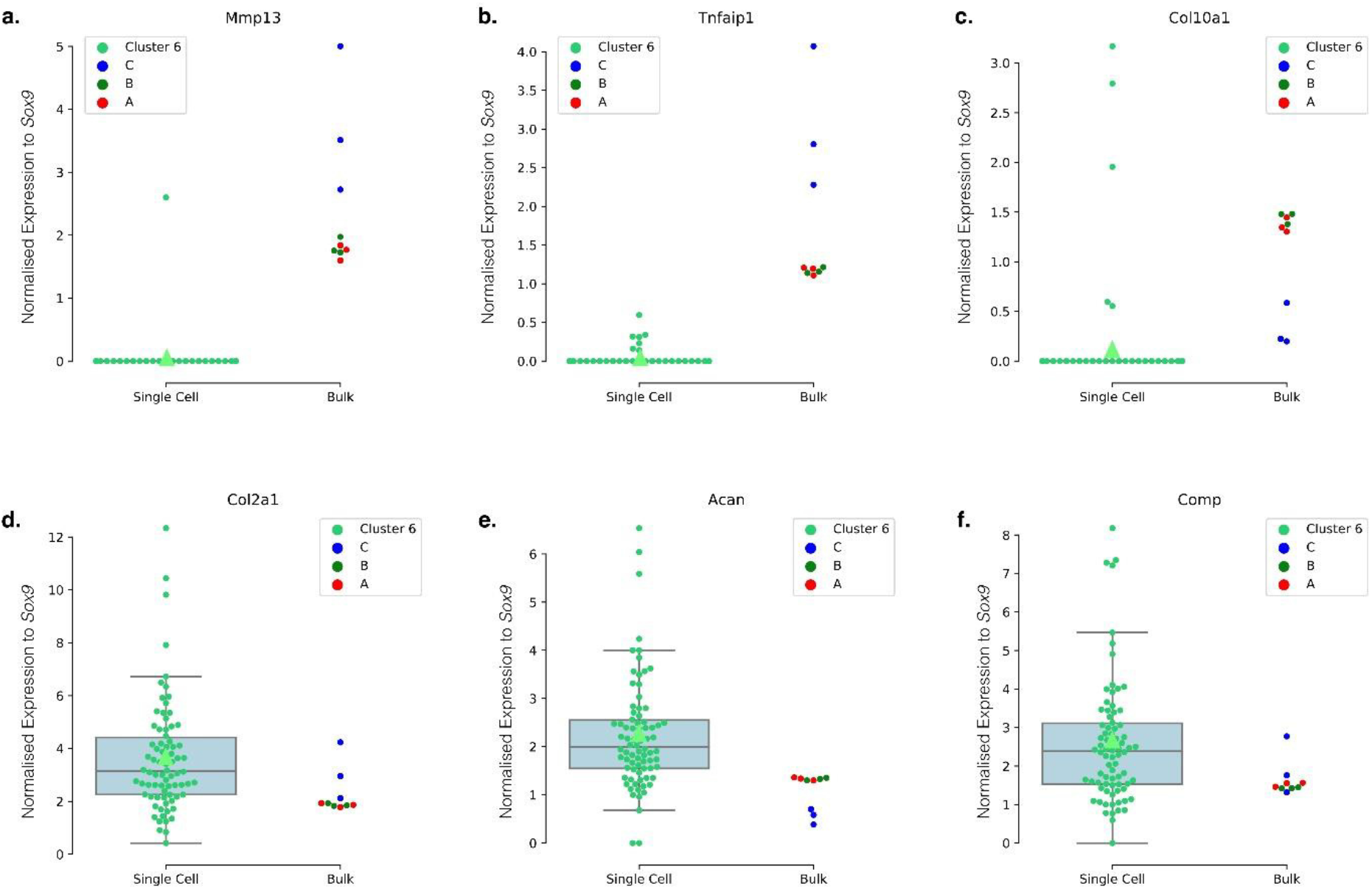
Normalized, specific gene expressions indicate no changes in transcriptome by digestions. **(a–f)**Boxplots of the Sox9-normalised expressions of the selected gene in cells of cluster 6. The mean expression in all cells is marked with a green triangle (scRNA-seq). The raw values are overlayed on the boxplot and coloured with a grey circle. The raw values of segments’ Sox9-normalised expression of the gene of interest (bulk RNA-seq). Segment A is shown in red; segment B is coloured in green, and segment C is coloured in blue. **(a)**Mmp13 **(b)**Tnfaip1 **(c)**Col10a1 **(d)**Col2a1 **(e)**Acan **(f)**Comp.

### Supplementary Results C: AC chondrocyte-specific genes identified by the intersection of bulk RNA-seq and scRNA-seq

## References

1. Kloppenburg, M. and F. Berenbaum, Osteoarthritis year in review 2019: epidemiology and therapy. Osteoarthritis Cartilage, 2020. 28(3): p. 242–248.

2. David, M.A., et al., Early, focal changes in cartilage cellularity and structure following surgically induced meniscal destabilization in the mouse. J Orthop Res, 2017. 35(3): p. 537–547.

3. Haase, T., et al., Discerning the spatio-temporal disease patterns of surgically induced OA mouse models. PLoS One, 2019. 14(4): p. e0213734.

4. Jeon, O.H., et al., Local clearance of senescent cells attenuates the development of post-traumatic osteoarthritis and creates a pro-regenerative environment. Nat Med, 2017. 23(6): p. 775–781.

5. Shalek, A.K. and M. Benson, Single-cell analyses to tailor treatments. Science translational medicine, 2017. 9(408): p. eaan4730.

6. Tang, F., et al., mRNA-Seq whole-transcriptome analysis of a single cell. Nat Methods, 2009. 6(5): p. 377–82.

7. Hayman, D.M., et al., The effects of isolation on chondrocyte gene expression. Tissue Eng, 2006. 12(9): p. 2573–81.

8. van den Brink, S.C., et al., Single-cell sequencing reveals dissociation-induced gene expression in tissue subpopulations. Nat Methods, 2017. 14(10): p. 935–936.

9. Fang, H. and F. Beier, Mouse models of osteoarthritis: modelling risk factors and assessing outcomes. Nat Rev Rheumatol, 2014. 10(7): p. 413–21.

10. Little, C.B. and D.J. Hunter, Post-traumatic osteoarthritis: from mouse models to clinical trials. Nat Rev Rheumatol, 2013. 9(8): p. 485–97.

11. Gardiner, M.D., et al., Transcriptional analysis of micro-dissected articular cartilage in post-traumatic murine osteoarthritis. Osteoarthritis Cartilage, 2015. 23(4): p. 616–28.

12. Archer, C.W. and P. Francis-West, The chondrocyte. Int J Biochem Cell Biol, 2003. 35(4): p. 401–4.

13. Aigner, T., et al., Histopathology atlas of animal model systems - overview of guiding principles. Osteoarthritis Cartilage, 2010. 18 Suppl 3: p. S2–6.

14. Hunziker, E.B., T.M. Quinn, and H.J. Hauselmann, Quantitative structural organization of normal adult human articular cartilage. Osteoarthritis Cartilage, 2002. 10(7): p. 564–72.

15. Chu, C.R., M. Szczodry, and S. Bruno, Animal models for cartilage regeneration and repair. Tissue Eng Part B Rev, 2010. 16(1): p. 105–15.

16. Malda, J., et al., Of mice, men and elephants: the relation between articular cartilage thickness and body mass. PLoS One, 2013. 8(2): p. e57683.

17. Kelly, N.H., N.P.T. Huynh, and F. Guilak, Single cell RNA-sequencing reveals cellular heterogeneity and trajectories of lineage specification during murine embryonic limb development. Matrix Biol, 2019.

18. Mizuhashi, K., et al., Growth Plate Borderline Chondrocytes Behave as Transient Mesenchymal Precursor Cells. J Bone Miner Res, 2019. 34(8): p. 1387–1392.

19. Baccin, C., et al., Combined single-cell and spatial transcriptomics reveal the molecular, cellular and spatial bone marrow niche organization. Nat Cell Biol, 2020. 22(1): p. 38–48.

20. Binch, A.L.A., et al., Class 3 semaphorins expression and association with innervation and angiogenesis within the degenerate human intervertebral disc. Oncotarget, 2015. 6(21): p. 18338–18354.

21. Liu, C.F., et al., SOX9 is dispensable for the initiation of epigenetic remodeling and the activation of marker genes at the onset of chondrogenesis. Development, 2018. 145(14).

22. Montagne, K., et al., High hydrostatic pressure induces pro-osteoarthritic changes in cartilage precursor cells: A transcriptome analysis. PLOS ONE, 2017. 12(8): p. e0183226.

23. Lau, D., et al., The cartilage-specific lectin C-type lectin domain family 3 member A (CLEC3A) enhances tissue plasminogen activator-mediated plasminogen activation. J Biol Chem, 2018. 293(1): p. 203–214.

24. Hu, S.I., et al., Human HtrA, an evolutionarily conserved serine protease identified as a differentially expressed gene product in osteoarthritic cartilage. J Biol Chem, 1998. 273(51): p. 34406–12.

25. Tsuchiya, A., et al., Expression of mouse HtrA1 serine protease in normal bone and cartilage and its upregulation in joint cartilage damaged by experimental arthritis. Bone, 2005. 37(3): p. 323–36.

26. Minchenko, O.H., et al., Effect of hypoxia on the expression of CCN2, PLAU, PLAUR, SLURP1, PLAT and ITGB1 genes in ERN1 knockdown U87 glioma cells. Ukr Biochem J, 2014. 86(4): p. 79–89.

27. Chang, J.C., et al., Global molecular changes in a tibial compression induced ACL rupture model of post-traumatic osteoarthritis. J Orthop Res, 2017. 35(3): p. 474–485.

28. Rapko, S., et al., Identification of the chondrocyte lineage using microfibril-associated glycoprotein-2, a novel marker that distinguishes chondrocytes from synovial cells. Tissue engineering. Part C, Methods, 2010. 16(6): p. 1367–1375.

29. Cote, A.J., et al., Single-cell differences in matrix gene expression do not predict matrix deposition. Nat Commun, 2016. 7: p. 10865.

30. Lepage, S.I.M., et al., Gene Expression Profile Is Different between Intact and Enzymatically Digested Equine Articular Cartilage. Cartilage, 2019: p. 1947603519833148.

31. Bensaude, O., Inhibiting eukaryotic transcription. Which compound to choose? How to evaluate its activity? Transcription, 2011. 2(3): p. 103–108.

32. Wu, Y.E., et al., Detecting Activated Cell Populations Using Single-Cell RNA-Seq. Neuron, 2017. 96(2): p. 313–329.e6.

33. Westendorf, K., et al., Unbiased transcriptomes of resting human CD4(+) CD45RO(+) T lymphocytes. Eur J Immunol, 2014. 44(6): p. 1866–9.

## References

1. Kim D, Pertea G, Trapnell C, Pimentel H, Kelley R, Salzberg SL. TopHat2: accurate alignment of transcriptomes in the presence of insertions, deletions and gene fusions. Genome Biol 2013; 14: R36.

2. Langmead B, Salzberg SL. Fast gapped-read alignment with Bowtie 2. Nature Methods 2012; 9: 357–359.

3. Liao Y, Smyth GK, Shi W. featureCounts: an efficient general purpose program for assigning sequence reads to genomic features. Bioinformatics 2014; 30: 923–930.

4. Love MI, Huber W, Anders S. Moderated estimation of fold change and dispersion for RNA-seq data with DESeq2. Genome Biology 2014; 15: 550.

5. Stuart T, Butler A, Hoffman P, Hafemeister C, Papalexi E, Mauck WM, et al. Comprehensive integration of single cell data. bioRxiv 2018: 460147.

6. Raudvere U, Kolberg L, Kuzmin I, Arak T, Adler P, Peterson H, et al. g:Profiler: a web server for functional enrichment analysis and conversions of gene lists (2019 update). Nucleic Acids Research 2019; 47: W191–W198.

## References

1. Chung J, Chen C, Paw BH. Heme metabolism and erythropoiesis. Current opinion in hematology 2012; 19: 156–162.

2. Hayman DM, Blumberg TJ, Scott CC, Athanasiou KA. The effects of isolation on chondrocyte gene expression. Tissue Eng 2006; 12: 2573–2581.

3. Mroczek S, Chlebowska J, Kuliński TM, Gewartowska O, Gruchota J, Cysewski D, et al. The non-canonical poly(A) polymerase FAM46C acts as an onco-suppressor in multiple myeloma. Nature Communications 2017; 8: 619.

4. Hu J-L, Liang H, Zhang H, Yang M-Z, Sun W, Zhang P, et al. FAM46B is a prokaryotic-like cytoplasmic poly(A) polymerase essential in human embryonic stem cells. Nucleic Acids Research 2020; 48: 2733–2748.

5. Huang X, Birk DE, Goetinck PF. Mice lacking matrilin-1 (cartilage matrix protein) have alterations in type II collagen fibrillogenesis and fibril organization. Dev Dyn 1999; 216: 434–441.

6. Nicolae C, Ko YP, Miosge N, Niehoff A, Studer D, Enggist L, et al. Abnormal collagen fibrils in cartilage of matrilin-1/matrilin-3-deficient mice. J Biol Chem 2007; 282: 22163–22175.

7. Binch ALA, Cole AA, Breakwell LM, Michael ALR, Chiverton N, Creemers LB, et al. Class 3 semaphorins expression and association with innervation and angiogenesis within the degenerate human intervertebral disc. Oncotarget 2015; 6: 18338–18354.

8. Catalano A. The neuroimmune semaphorin-3A reduces inflammation and progression of experimental autoimmune arthritis. J Immunol 2010; 185: 6373–6383.

9. Liu CF, Angelozzi M, Haseeb A, Lefebvre V. SOX9 is dispensable for the initiation of epigenetic remodeling and the activation of marker genes at the onset of chondrogenesis. Development 2018; 145.

10. Zhu S, Qiu H, Bennett S, Kuek V, Rosen V, Xu H, et al. Chondromodulin-1 in health, osteoarthritis, cancer, and heart disease. Cellular and Molecular Life Sciences 2019; 76: 4493–4502.

11. Aigner T, Fundel K, Saas J, Gebhard PM, Haag J, Weiss T, et al. Large-scale gene expression profiling reveals major pathogenetic pathways of cartilage degeneration in osteoarthritis. Arthritis & Rheumatism 2006; 54: 3533–3544.

12. Regan E, Flannelly J, Bowler R, Tran K, Nicks M, Carbone BD, et al. Extracellular superoxide dismutase and oxidant damage in osteoarthritis. Arthritis & Rheumatism 2005; 52: 3479–3491.

13. Shi Y, Hu X, Cheng J, Zhang X, Zhao F, Shi W, et al. A small molecule promotes cartilage extracellular matrix generation and inhibits osteoarthritis development. Nature Communications 2019; 10: 1914.

14. Qin S, Ingle JN, Liu M, Yu J, Wickerham DL, Kubo M, et al. Calmodulin-like protein 3 is an estrogen receptor alpha coregulator for gene expression and drug response in a SNP, estrogen, and SERM-dependent fashion. Breast cancer research : BCR 2017; 19: 95–95.

15. Ashwell MS, O’Nan AT, Gonda MG, Mente PL. Gene expression profiling of chondrocytes from a porcine impact injury model. Osteoarthritis and Cartilage 2008; 16: 936–946.

16. Blazek AD, Nam J, Gupta R, Pradhan M, Perera P, Weisleder NL, et al. Exercise-driven metabolic pathways in healthy cartilage. Osteoarthritis and cartilage 2016; 24: 1210–1222.

17. Budde B, Blumbach K, Ylostalo J, Zaucke F, Ehlen HW, Wagener R, et al. Altered integration of matrilin-3 into cartilage extracellular matrix in the absence of collagen IX. Mol Cell Biol 2005; 25: 10465–10478.

18. Opolka A, Ratzinger S, Schubert T, Spiegel HU, Grifka J, Bruckner P, et al. Collagen IX is indispensable for timely maturation of cartilage during fracture repair in mice. Matrix Biol 2007; 26: 85–95.

19. Jenner F, Ijpma A, Cleary M, Heijsman D, Narcisi R, van der Spek PJ, et al. Differential gene expression of the intermediate and outer interzone layers of developing articular cartilage in murine embryos. Stem cells and development 2014; 23: 1883–1898.

20. Luo X, Wang J, Wei X, Wang S, Wang A. Knockdown of lncRNA MFI2-AS1 inhibits lipopolysaccharide-induced osteoarthritis progression by miR-130a-3p/TCF4. Life Sciences 2020; 240: 117019.

21. St-Jacques B, Hammerschmidt M, McMahon AP. Indian hedgehog signaling regulates proliferation and differentiation of chondrocytes and is essential for bone formation. Genes & development 1999; 13: 2072–2086.

22. Yang J, Andre P, Ye L, Yang Y-Z. The Hedgehog signalling pathway in bone formation. International Journal of Oral Science 2015; 7: 73–79.

23. Lui JCK, Andrade AC, Forcinito P, Hegde A, Chen W, Baron J, et al. Spatial and temporal regulation of gene expression in the mammalian growth plate. Bone 2010; 46: 1380–1390.

24. Monterrat C, Boal F, Grise F, Hemar A, Lang J. Synaptotagmin 8 is expressed both as a calcium-insensitive soluble and membrane protein in neurons, neuroendocrine and endocrine cells. Biochim Biophys Acta 2006; 1763: 73–81.

25. Mori Y, Chung UI, Tanaka S, Saito T. Determination of differential gene expression profiles in superficial and deeper zones of mature rat articular cartilage using RNA sequencing of laser microdissected tissue specimens. Biomed Res 2014; 35: 263–270.

26. Seuffert F, Weidner D, Baum W, Schett G, Stock M. Upper zone of growth plate and cartilage matrix associated protein protects cartilage during inflammatory arthritis. Arthritis Research & Therapy 2018; 20: 88.

27. Lee Y. The role of interleukin-17 in bone metabolism and inflammatory skeletal diseases. BMB reports 2013; 46: 479–483.

28. Moseley TA, Haudenschild DR, Rose L, Reddi AH. Interleukin-17 family and IL-17 receptors. Cytokine Growth Factor Rev 2003; 14: 155–174.

29. Lorenzo P, Bayliss MT, Heinegard D. A novel cartilage protein (CILP) present in the mid-zone of human articular cartilage increases with age. J Biol Chem 1998; 273: 23463–23468.

30. Lorenzo P, Bayliss MT, Heinegård D. Altered patterns and synthesis of extracellular matrix macromolecules in early osteoarthritis. Matrix Biology 2004; 23: 381–391.

31. Seki S, Kawaguchi Y, Chiba K, Mikami Y, Kizawa H, Oya T, et al. A functional SNP in CILP, encoding cartilage intermediate layer protein, is associated with susceptibility to lumbar disc disease. Nat Genet 2005; 37: 607–612.

32. Gardiner MD, Vincent TL, Driscoll C, Burleigh A, Bou-Gharios G, Saklatvala J, et al. Transcriptional analysis of micro-dissected articular cartilage in post-traumatic murine osteoarthritis. Osteoarthritis Cartilage 2015; 23: 616–628.

33. Montagne K, Onuma Y, Ito Y, Aiki Y, Furukawa KS, Ushida T. High hydrostatic pressure induces pro-osteoarthritic changes in cartilage precursor cells: A transcriptome analysis. PLOS ONE 2017; 12: e0183226.

34. Ramos YF, Bos SD, van der Breggen R, Kloppenburg M, Ye K, Lameijer EW, et al. A gain of function mutation in TNFRSF11B encoding osteoprotegerin causes osteoarthritis with chondrocalcinosis. Ann Rheum Dis 2015; 74: 1756–1762.

35. Ruiz AR, Houtman, E., Van Hoolwerff, M., Lakenberg, N., Nelissen, R.G., Meulenbelt, I., Ramos, Y.F. Towards the elucidation of the role of osteoprotegerin in the development of osteoarthritis. OARSI World Congress, vol. 26: Osteoarthritis & Cartilage 2018:S87.

36. Arnott JA, Lambi AG, Mundy C, Hendesi H, Pixley RA, Owen TA, et al. The role of connective tissue growth factor (CTGF/CCN2) in skeletogenesis. Critical reviews in eukaryotic gene expression 2011; 21: 43–69.

37. Omoto S, Nishida K, Yamaai Y, Shibahara M, Nishida T, Doi T, et al. Expression and localization of connective tissue growth factor (CTGF/Hcs24/CCN2) in osteoarthritic cartilage. Osteoarthritis Cartilage 2004; 12: 771–778.

38. Moon P, Penuela S, Laird DW, Beier F. Pannexin 3, but not pannexin 1 is an important pro-catabolic mediator in osteoarthritis. Osteoarthritis and Cartilage 2016; 24: S153.

39. Moon PM, Penuela S, Barr K, Khan S, Pin CL, Welch I, et al. Deletion of Panx3 Prevents the Development of Surgically Induced Osteoarthritis. Journal of molecular medicine (Berlin, Germany) 2015; 93: 845–856.

40. Lories RJ, Peeters J, Bakker A, Tylzanowski P, Derese I, Schrooten J, et al. Articular cartilage and biomechanical properties of the long bones in Frzb-knockout mice. Arthritis Rheum 2007; 56: 4095–4103.

41. Thysen S, Cailotto F, Luyten FP, Lories RJ. FRZB is a critical modulator of canonical WNT signalling in cartilage biology. Osteoarthritis and Cartilage 2013; 21: S112.

42. White AH, Watson RE, Newman B, Freemont AJ, Wallis GA. Annexin VIII is differentially expressed by chondrocytes in the mammalian growth plate during endochondral ossification and in osteoarthritic cartilage. J Bone Miner Res 2002; 17: 1851–1858.

43. Blaydon DC, Ishii Y, O’Toole EA, Unsworth HC, Teh MT, Ruschendorf F, et al. The gene encoding R-spondin 4 (RSPO4), a secreted protein implicated in Wnt signaling, is mutated in inherited anonychia. Nat Genet 2006; 38: 1245–1247.

44. Henry SP, Liang S, Akdemir KC, de Crombrugghe B. The postnatal role of Sox9 in cartilage. Journal of bone and mineral research : the official journal of the American Society for Bone and Mineral Research 2012; 27: 2511–2525.

45. Kronke G, Uderhardt S, Kim KA, Stock M, Scholtysek C, Zaiss MM, et al. R-spondin 1 protects against inflammatory bone damage during murine arthritis by modulating the Wnt pathway. Arthritis Rheum 2010; 62: 2303–2312.

46. Zheng J, Wu C, Ma W, Zhang Y, Hou T, Xu H, et al. Abnormal expression of chondroitin sulphate N-acetylgalactosaminyltransferase 1 and Hapln-1 in cartilage with Kashin– Beck disease and primary osteoarthritis. International Orthopaedics 2013; 37: 2051–2059.

47. Mao JR, Taylor G, Dean WB, Wagner DR, Afzal V, Lotz JC, et al. Tenascin-X deficiency mimics Ehlers-Danlos syndrome in mice through alteration of collagen deposition. Nat Genet 2002; 30: 421–425.

48. Tillgren V, Ho JC, Onnerfjord P, Kalamajski S. The novel small leucine-rich protein chondroadherin-like (CHADL) is expressed in cartilage and modulates chondrocyte differentiation. J Biol Chem 2015; 290: 918–925.

49. Kiepe D, Ulinski T, Powell DR, Durham SK, Mehls O, Tönshoff B. Differential effects of insulin-like growth factor binding proteins-1, -2, -3, and -6 on cultured growth plate chondrocytes. Kidney International 2002; 62: 1591–1600.

50. Turlo AJ, Mueller-Breckenridge AJ, Zamboulis DE, Tew SR, Canty-Laird EG, Clegg PD. Insulin-like growth factor binding protein (IGFBP6) is a cross-species tendon marker. Eur Cell Mater 2019; 38: 123–136.

51. Zhang L, Smith DW, Gardiner BS, Grodzinsky AJ. Modeling the Insulin-Like Growth Factor System in Articular Cartilage. PloS one 2013; 8: e66870–e66870.

52. Aki T, Hashimoto K, Ogasawara M, Itoi E. A whole-genome transcriptome analysis of articular chondrocytes in secondary osteoarthritis of the hip. PloS one 2018; 13: e0199734–e0199734.

53. Su W, Xie W, Shang Q, Su B. The Long Noncoding RNA MEG3 Is Downregulated and Inversely Associated with VEGF Levels in Osteoarthritis. Biomed Res Int 2015; 2015: 356893.

54. Wang A, Hu N, Zhang Y, Chen Y, Su C, Lv Y, et al. MEG3 promotes proliferation and inhibits apoptosis in osteoarthritis chondrocytes by miR-361-5p/FOXO1 axis. BMC Medical Genomics 2019; 12: 201.

55. Xu J, Xu Y. The lncRNA MEG3 downregulation leads to osteoarthritis progression via miR-16/SMAD7 axis. Cell & bioscience 2017; 7: 69–69.

56. Jamil NS, Azfer A, Sarah HE, Salter DM. NG2/CSPG4 regulates aggrecanase and MMP expression in human chondrocytes. Osteoarthritis and Cartilage 2012; 20: S44.

57. Stallcup WB. The NG2 proteoglycan: past insights and future prospects. J Neurocytol 2002; 31: 423–435.

58. Lau D, Elezagic D, Hermes G, Morgelin M, Wohl AP, Koch M, et al. The cartilage-specific lectin C-type lectin domain family 3 member A (CLEC3A) enhances tissue plasminogen activator-mediated plasminogen activation. J Biol Chem 2018; 293: 203–214.

59. Acosta E, Avila J, Mobasheri A, Martin-Vasallo P. Na+, K+-ATPase genes are down-regulated during adipose stem cell differentiation. Front Biosci (Elite Ed) 2011; 3: 1229–1240.

60. Mobasheri A, Trujillo E, Arteaga MF, Martin-Vasallo P. Na(+), K(+)-ATPase subunit composition in a human chondrocyte cell line; evidence for the presence of alpha1, alpha3, beta1, beta2 and beta3 isoforms. Int J Mol Sci 2012; 13: 5019–5034.

61. Hu SI, Carozza M, Klein M, Nantermet P, Luk D, Crowl RM. Human HtrA, an evolutionarily conserved serine protease identified as a differentially expressed gene product in osteoarthritic cartilage. J Biol Chem 1998; 273: 34406–34412.

62. Tsuchiya A, Yano M, Tocharus J, Kojima H, Fukumoto M, Kawaichi M, et al. Expression of mouse HtrA1 serine protease in normal bone and cartilage and its upregulation in joint cartilage damaged by experimental arthritis. Bone 2005; 37: 323–336.

63. Johnson HJ, Rosenberg L, Choi HU, Garza S, Hook M, Neame PJ. Characterization of epiphycan, a small proteoglycan with a leucine-rich repeat core protein. J Biol Chem 1997; 272: 18709–18717.

64. Nuka S, Zhou W, Henry SP, Gendron CM, Schultz JB, Shinomura T, et al. Phenotypic characterization of epiphycan-deficient and epiphycan/biglycan double-deficient mice. Osteoarthritis and cartilage 2010; 18: 88–96.

65. Steiglitz BM, Keene DR, Greenspan DS. PCOLCE2 encodes a functional procollagen C-proteinase enhancer (PCPE2) that is a collagen-binding protein differing in distribution of expression and post-translational modification from the previously described PCPE1. J Biol Chem 2002; 277: 49820–49830.

66. Song EK, Jeon J, Jang DG, Kim HE, Sim HJ, Kwon KY, et al. ITGBL1 modulates integrin activity to promote cartilage formation and protect against arthritis. Sci Transl Med 2018; 10.

67. Hatfield JT, Anderson PJ, Powell BC. Retinol-binding protein 4 is expressed in chondrocytes of developing mouse long bones: implications for a local role in formation of the secondary ossification center. Histochem Cell Biol 2013; 139: 727–734.

68. Scotece M, Koskinen-Kolasa A, Pemmari A, Leppanen T, Hamalainen M, Moilanen T, et al. Novel adipokine associated with OA: retinol binding protein 4 (RBP4) is produced by cartilage and is correlated with MMPs in osteoarthritis patients. Inflamm Res 2020; 69: 415–421.

69. Schubert T, Schlegel J, Schmid R, Opolka A, Grässel S, Humphries M, et al. Modulation of cartilage differentiation by melanoma inhibiting activity/cartilage-derived retinoic acid-sensitive protein (MIA/CD-RAP). Experimental & Molecular Medicine 2010; 42: 166–174.

70. Zhang Y, Morgan BJ, Smith R, Fellows CR, Thornton C, Snow M, et al. Platelet-rich plasma induces post-natal maturation of immature articular cartilage and correlates with LOXL1 activation. Scientific reports 2017; 7: 3699–3699.

71. Hankenson KD, Hormuzdi SG, Meganck JA, Bornstein P. Mice with a disruption of the thrombospondin 3 gene differ in geometric and biomechanical properties of bone and have accelerated development of the femoral head. Molecular and cellular biology 2005; 25: 5599–5606.

72. Jeschke A, Bonitz M, Simon M, Peters S, Baum W, Schett G, et al. Deficiency of Thrombospondin-4 in Mice Does Not Affect Skeletal Growth or Bone Mass Acquisition, but Causes a Transient Reduction of Articular Cartilage Thickness. PLoS One 2015; 10: e0144272.

73. Maly K, Schaible I, Riegger J, Brenner RE, Meurer A, Zaucke F. The Expression of Thrombospondin-4 Correlates with Disease Severity in Osteoarthritic Knee Cartilage. International journal of molecular sciences 2019; 20: 447.

74. Gerhardt B, Leesman L, Burra K, Snowball J, Rosenzweig R, Guzman N, et al. Notum attenuates Wnt/beta-catenin signaling to promote tracheal cartilage patterning. Dev Biol 2018; 436: 14–27.

75. Ikeuchi T, de Vega S, Forcinito P, Doyle AD, Amaral J, Rodriguez IR, et al. Extracellular Protein Fibulin-7 and Its C-Terminal Fragment Have In Vivo Antiangiogenic Activity. Scientific Reports 2018; 8: 17654.

76. Janune D, Abd El Kader T, Aoyama E, Nishida T, Tabata Y, Kubota S, et al. Novel role of CCN3 that maintains the differentiated phenotype of articular cartilage. J Bone Miner Metab 2017; 35: 582–597.

77. Rapko S, Zhang M, Richards B, Hutto E, Dethlefsen S, Duguay S. Identification of the chondrocyte lineage using microfibril-associated glycoprotein-2, a novel marker that distinguishes chondrocytes from synovial cells. Tissue engineering. Part C, Methods 2010; 16: 1367–1375.

78. Geister KA, Lopez-Jimenez AJ, Houghtaling S, Ho TH, Vanacore R, Beier DR. Loss of function of Colgalt1 disrupts collagen post-translational modification and causes musculoskeletal defects. Dis Model Mech 2019; 12.

79. Chang JC, Sebastian A, Murugesh DK, Hatsell S, Economides AN, Christiansen BA, et al. Global molecular changes in a tibial compression induced ACL rupture model of post-traumatic osteoarthritis. J Orthop Res 2017; 35: 474–485.

80. Rai MF, Sandell LJ, Zhang B, Wright RW, Brophy RH. RNA Microarray Analysis of Macroscopically Normal Articular Cartilage from Knees Undergoing Partial Medial Meniscectomy: Potential Prediction of the Risk for Developing Osteoarthritis. PloS one 2016; 11: e0155373–e0155373.

81. Jeon J, Oh H, Lee G, Ryu JH, Rhee J, Kim JH, et al. Cytokine-like 1 knock-out mice (Cytl1−/−) show normal cartilage and bone development but exhibit augmented osteoarthritic cartilage destruction. J Biol Chem 2011; 286: 27206–27213.

82. Zhu S, Kuek V, Bennett S, Xu H, Rosen V, Xu J. Protein Cytl1: its role in chondrogenesis, cartilage homeostasis, and disease. Cellular and Molecular Life Sciences 2019; 76: 3515–3523.

83. Maeda T, Jikko A, Abe M, Yokohama-Tamaki T, Akiyama H, Furukawa S, et al. Cartducin, a paralog of Acrp30/adiponectin, is induced during chondrogenic differentiation and promotes proliferation of chondrogenic precursors and chondrocytes. Journal of Cellular Physiology 2006; 206: 537–544.

84. Murayama MA, Kakuta S, Maruhashi T, Shimizu K, Seno A, Kubo S, et al. CTRP3 plays an important role in the development of collagen-induced arthritis in mice. Biochemical and Biophysical Research Communications 2014; 443: 42–48.

85. Robra L, Thirumalai V. The Intracellular Signaling Molecule Darpp-32 Is a Marker for Principal Neurons in the Cerebellum and Cerebellum-Like Circuits of Zebrafish. Frontiers in neuroanatomy 2016; 10: 81–81.

86. Woods A, James CG, Wang G, Dupuis H, Beier F. Control of chondrocyte gene expression by actin dynamics: a novel role of cholesterol/Ror-alpha signalling in endochondral bone growth. Journal of cellular and molecular medicine 2009; 13: 3497–3516.

87. Minchenko OH, Kharkova AP, Kubaichuk KI, Minchenko DO, Hlushchak NA, Kovalevska OV. Effect of hypoxia on the expression of CCN2, PLAU, PLAUR, SLURP1, PLAT and ITGB1 genes in ERN1 knockdown U87 glioma cells. Ukr Biochem J 2014; 86: 79–89.

88. Schultz M, Jin W, Waheed A, Moed BR, Sly W, Zhang Z. Expression profile of carbonic anhydrases in articular cartilage. Histochem Cell Biol 2011; 136: 145–151.

89. Schmid R, Schiffner S, Opolka A, Grässel S, Schubert T, Moser M, et al. Enhanced cartilage regeneration in MIA/CD-RAP deficient mice. Cell Death & Disease 2010; 1: e97–e97.

90. Tscheudschilsuren G, Bosserhoff AK, Schlegel J, Vollmer D, Anton A, Alt V, et al. Regulation of mesenchymal stem cell and chondrocyte differentiation by MIA. Exp Cell Res 2006; 312: 63–72.

91. Koltes JE, Kumar D, Kataria RS, Cooper V, Reecy JM. Transcriptional profiling of PRKG2-null growth plate identifies putative down-stream targets of PRKG2. BMC research notes 2015; 8: 177–177.

92. Hessle L, Stordalen GA, Wenglén C, Petzold C, Tanner EK, Brorson S-H, et al. The Skeletal Phenotype of Chondroadherin Deficient Mice. PLOS ONE 2013; 8: e63080.

93. Gari MA, AlKaff M, Alsehli HS, Dallol A, Gari A, Abu-Elmagd M, et al. Identification of novel genetic variations affecting osteoarthritis patients. BMC medical genetics 2016; 17: 68–68.

94. Chan DD, Xiao WF, Li J, de la Motte CA, Sandy JD, Plaas A. Deficiency of hyaluronan synthase 1 (Has1) results in chronic joint inflammation and widespread intra-articular fibrosis in a murine model of knee joint cartilage damage. Osteoarthritis and cartilage 2015; 23: 1879–1889.

95. Siiskonen H, Oikari S, Pasonen-Seppänen S, Rilla K. Hyaluronan Synthase 1: A Mysterious Enzyme with Unexpected Functions. Frontiers in Immunology 2015; 6.

